# (p)ppGpp-mediated GTP homeostasis ensures the survival and antibiotic tolerance of *Staphylococcus aureus* during starvation by preserving the proton motive force

**DOI:** 10.1101/2024.02.06.579068

**Authors:** Andrea Salzer, Sophia Ingrassia, Lisa Sauer, Johanna Rapp, Ronja Dobritz, Jennifer Müller, Hannes Link, Christiane Wolz

## Abstract

Upon nutrient limitation bacteria enter a nongrowing state, which allow bacterial survival and antibiotic tolerance. The mechanisms whether and how the messenger molecule (p)ppGpp contributes to the transition in Firmicutes is debated. Here we show for *Staphylococcus aureus* that (p)ppGpp-dependent restriction of the GTP pool is essential for the culturability of starved cells and for antibiotic tolerance. Elevated GTP levels in a starving (p)ppGpp-deficient mutant lead to a division-incompetent, dormant state characterized by reduced metabolic activity and alterations in membrane function and architecture. GTP level control of nucleotide sensitive promoters result in transcriptional downregulation of gene of the TCA cycle and electron transport chain. Increasing transcription of *qoxABCD*, a terminal oxidase of the respiratory chain, through mutation of the transcriptional start site partially restored the culturability of the (p)ppGpp-deficient mutant. Furthermore, we showed that the maintenance of proton motive force under nutritional stress contributes to antibiotic tolerance, supporting the idea of applying (p)ppGpp or PMF inhibitors to combat antibiotic-tolerant bacteria.

## Introduction

Bacteria need to switch between rapid growth under favourable conditions and a nongrowing but stress-tolerant state under limiting conditions. In pathogenic bacteria, such a nongrowing state is often associated with persistent infections and antibiotic tolerance ^1–3^. Nongrowing bacteria continue to synthesize macromolecules, have a basal metabolism ^4^ and are prepared to resume growth. Recent studies have demonstrated that the messenger molecule (p)ppGpp is central to coordinating optimal resource allocation under nutrient-limited conditions ^5^. For *Escherichia coli*, (p)ppGpp was postulated to act as the major regulator of growth rate control ^6,7^. Even subtle fluctuations in levels of (p)ppGpp or basal levels that might be below the detection limit, such as during entry into the stationary growth phase, can modulate cellular processes and growth rate ^8^. (p)ppGpp specifically interferes with replication, translation, and transcription ^9^. In proteobacteria, (p)ppGpp directly binds to RNA polymerase to regulate transcription. In Firmicutes, however, (p)ppGpp does not interfere with RNA polymerase activity ^10^. In these organisms, (p)ppGpp regulate transcription mainly by controlling intracellular GTP levels ^11–14^. GTP homeostasis is controlled by specific inhibitors of enzymes involved in GTP synthesis, including the guanylate kinase Gmk, the IMP dehydrogenase GuaB, the hypoxanthine-guanine phosphoribosyltransferase HrpT, and the transcriptional regulator PurR ^15^. Thus, (p)ppGpp synthesis correlates inversely with the GTP pool. A decrease in GTP levels during the stringent response may affect transcription via the GTP-sensitive transcriptional regulator CodY ^16^ or through regulation of initiation nucleotide (iNTP)-sensitive promoters ^11,14,17^.

The role of (p)ppGpp in stationary phase survival and bacterial tolerance in *Staphylococcus aureus* is controversial ^3,18–20^. In this human pathogen and other Firmicutes species, synthesis and degradation of the messenger (p)ppGpp are orchestrated by the bifunctional Rel enzyme with synthetase and hydrolase activity and by the two synthetases SasA/RelP and SasB/RelQ ^9,15^. Induction of the stringent response via amino acid limitation (Rel) or cell wall stress (RelPQ) contributes to pathogenicity and antibiotic tolerance ^12,20^. However, the role of basal levels of (p)ppGpp in bacterial survival under nonstressed conditions is less clear. Here, we show that under nutrient stress, (p)ppGpp is essential for maintaining GTP homeostasis and thereby preserving electron transport chain activity and the proton motive force (PMF), which ensures culturability and antibiotic tolerance.

## Results

### (p)ppGpp contributes to culturability in the stationary phase

To study the requirement of (p)ppGpp synthesis for stationary phase survival, we monitored the growth and culturability (determined by CFU/ml enumeration) of *S. aureus* wildtype strain HG001 and its isogenic (p)ppGpp^0^ mutant in chemically defined medium (CDM) for 24 h. (p)ppGpp^0^ is unable to synthesize (p)ppGpp due to mutations in all three (p)ppGpp synthetases (full deletion of *rel*, synthetase mutations in *relP* and *relQ)* ^21^. There was no significant difference in growth measured by optical density (OD_600_) between the two strains (Fig. 1A). However, after entry into the stationary phase, the culturability of the (p)ppGpp^0^ mutant quickly started to decrease until an approximately one log reduction in CFU/ml was observed after 24 h compared to that of wildtype (Figs. 1B & S1A). Reduced culturability of the (p)ppGpp^0^ mutant was also observed in the background of the USA300 JE2 or SH1000 strains (the SigB-positive derivative and phage-cured derivative of HG001, respectively) (Fig. 1C). We next induced Rel-dependent (p)ppGpp synthesis in exponentially growing bacteria with a subinhibitory concentration of mupirocin, a tRNA synthetase inhibitor. These conditions mimicked amino acid starvation and resulted in significantly reduced culturability of the (p)ppGpp^0^ mutant (Fig. S1B). Furthermore, we monitored expression levels of the known stringent response-activated genes *psmα* and *rsaD* ^22,23^ during the mid- and late stationary phases (17 h and 24 h) in wildtype and (p)ppGpp^0^. We observed a stringent response-like expression pattern for *psmα* and strong derepression of *rsaD* in wildtype, indicating elevated (p)ppGpp levels (Fig. S1C). As expected, in the (p)ppGpp^0^ mutant, expression of stringent response genes was not induced, but *rsaD* expression was strongly downregulated, suggesting elevated intracellular GTP levels (Fig. S1C). Individual complementation of (p)ppGpp^0^ with *rel_syn_*, *relP* or *relQ* (Fig. 1D) confirmed that Rel and RelQ contribute to basal (p)ppGpp levels to ensure survival during the stationary phase. Thus, (p)ppGpp is essential for bacterial culturability in the stationary phase as well as under induced stress conditions in exponentially growing bacteria.

**Fig. 1.**
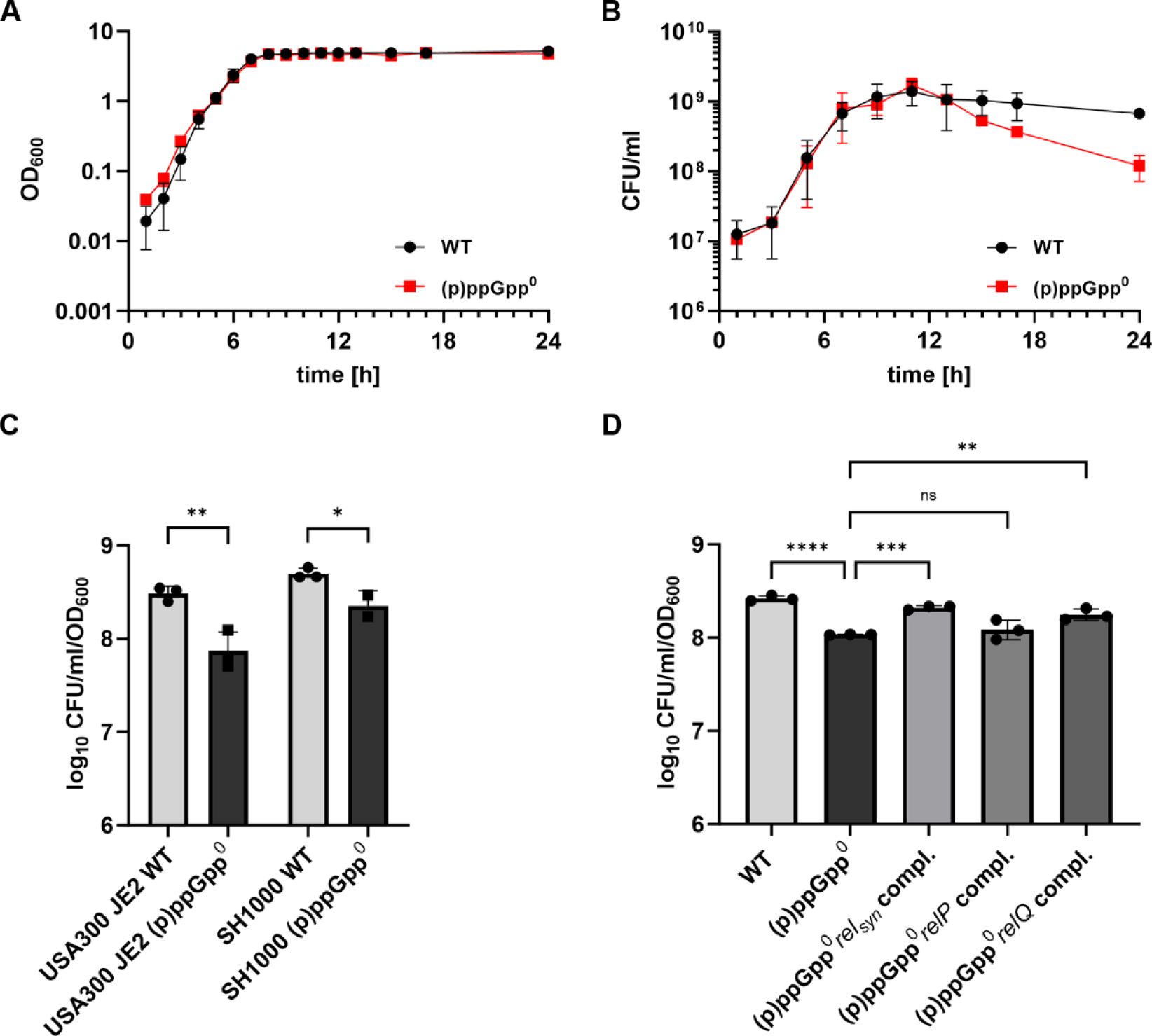
Culturability of the (p)ppGpp^0^ mutant decreases during the stationary growth phase. (A) *S. aureus* HG001 wildtype (WT) and the isogenic (p)ppGpp^0^ strain were cultured in CDM ^47^ and growth was monitored by optical density OD_600_ and (B) CFU/ml determination. Data are shown as mean ± SD (n=3 biological replicates). (C) USA300 JE2 and SH1000 and their isogenic (p)ppGpp^0^ mutants were grown in CDM for 24h and culturability was determined by CFU/ml enumeration normalized to OD_600_. Data are shown as mean ± SD (n=3 biological replicates). Statistical significance was determined by two-tailed unpaired t-tests performed on log_10_ transformed data (**p-value <0.01, *p-value <0.05). (D) Strains were grown for 24h in CDM and culturability was determined by CFU/ml enumeration normalized to OD_600_. Data shown are mean ± SD (n=3 biological replicates). Statistical significance was determined by one-way analysis of variance (ANOVA) with a Šidák’s post-test on log_10_ transformed data (****p-value <0.0001, ***p-value <0.001, **p-value <0.01, ns >0.05).

### (p)ppGpp-dependent GTP homeostasis is essential for stationary phase culturability independent of CodY

(p)ppGpp has been shown to play an important role in controlling GTP homeostasis ^13,14^. Therefore, we assessed the viability of different mutants in which GTP metabolism was disrupted after growth to the late stationary phase. The pleiotropic repressor CodY, when loaded with GTP, inhibits expression of several biosynthesis-related gene clusters and virulence genes. CodY target genes are derepressed under stringent conditions in a (p)ppGpp/lowGTP-dependent manner ^20^. We therefore hypothesized that the decreased survival of the (p)ppGpp^0^ mutant might be ascribed to diminished expression of CodY target genes. However, deletion of *codY* did not increase the culturability of the (p)ppGpp^0^ mutant (Fig. 2B). PurR is a repressor of the purine operon ^24,25^, and in *B. subtilis*, it is allosterically activated via (p)ppGpp ^15^ (Fig. 2A). Thus, deletion of *purR* in the (p)ppGpp^0^ mutant will likely lead to increased GTP levels. This mutant was even less culturable in the stationary phase, indicating that elevated GTP levels are detrimental to bacterial culturability (Fig. 2B). Accordingly, lowering the GTP pool should promote bacterial survival. Indeed, deletion of the GTP synthesis operon *guaBA* rescued the defect in the (p)ppGpp^0^ mutant (Fig. 2B). We next manipulated levels of GTP via supplementation with guanine, which feeds into the salvage pathway of GTP synthesis (Fig. 2A). This resulted in a dose-dependent decrease in the culturability of the (p)ppGpp^0^ mutant (Fig. 2C).

**Fig. 2.**
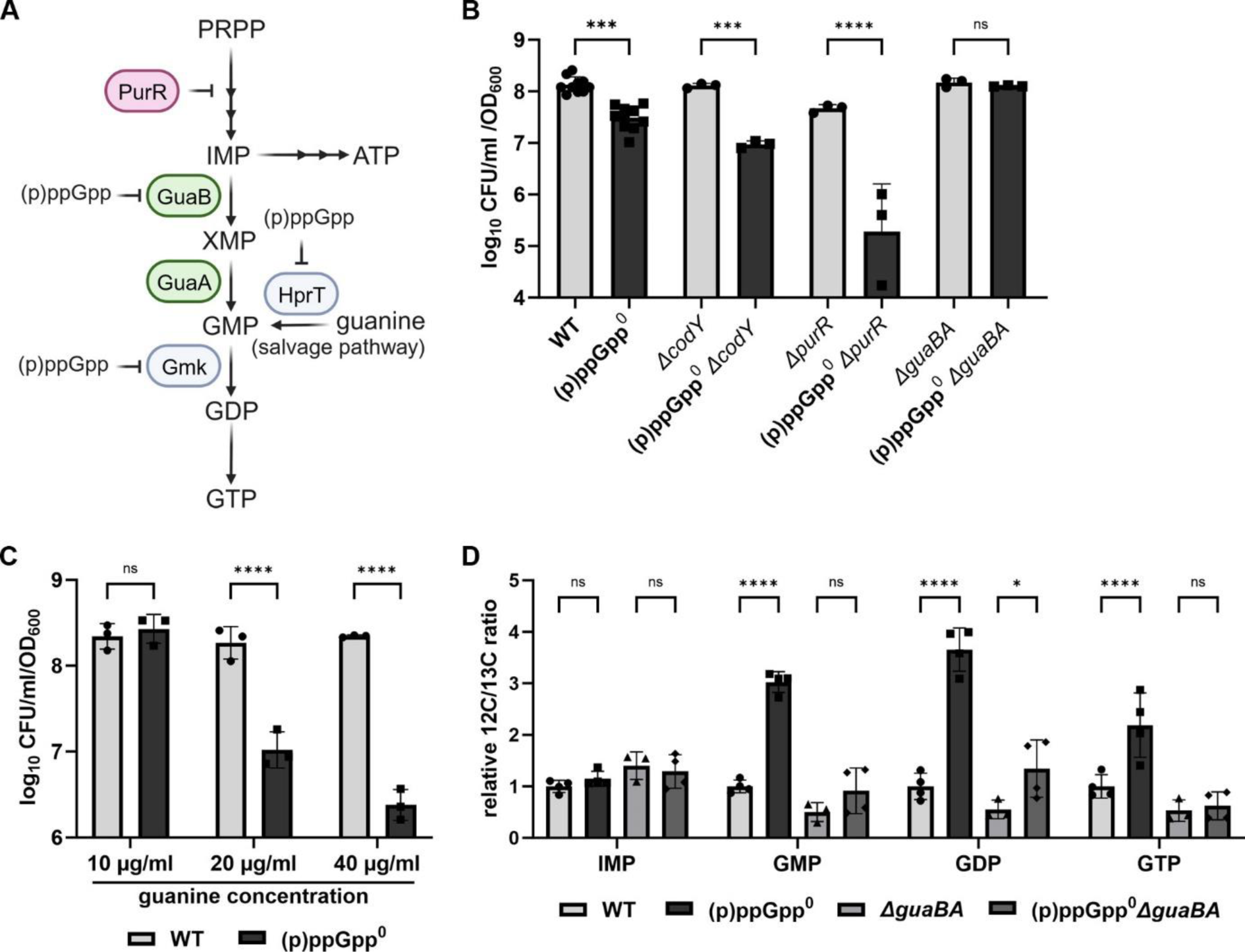
(p)ppGpp is essential for GTP homeostasis during stationary phase starvation. (A) The purine biosynthesis pathway is regulated by PurR repression and by (p)ppGpp repressing the biosynthetic enzymes GuaB, HrpT and Gmk. (B) Strains were grown in CDM for 24h and survival was determined by CFU/ml enumeration normalized to OD_600_. Data shown are mean ± SD (n≥3). Statistical significance was determined by one-way analysis of variance (ANOVA) with a Tukey’s post-test on log_10_ transformed data (****p-value <0.0001, ***p-value <0.001, ns >0.05). (C) HG001 wildtype (WT) and (p)ppGpp^0^ were grown in CDM with varying concentrations of guanine for 24h and culturability was determined by CFU/ml enumeration normalized to OD_600_. Data shown are mean ± SD (n=3). Statistical significance was determined by two-way way analysis of variance (ANOVA) with a Šidák post-test (****p-value <0.0001, ns >0.05). (D) Relative nucleotide levels were determined from late stationary phase cells by LC-MS/MS and normalized to OD_600_. Data shown are mean ± SD (n=4). Statistical significance was determined by two-way analysis of variance (ANOVA) with a Tukey’s post-test (****p-value <0.0001, ***p-value <0.001, **p-value <0.01, ns >0.05).

We used LC–MS/MS analysis to confirm the anticipated nucleotide levels in the (p)ppGpp^0^ and *ΔguaAB* mutants. As expected, levels of all guanine nucleotides (GMP, GDP and GTP) were significantly elevated in the (p)ppGpp^0^ mutant in the late stationary phase (Fig. 2D). Lower levels were observed in wildtype, likely due to (p)ppGpp-dependent inhibition of biosynthetic enzymes, including GuaB (Fig. 2A). Consistently, deletion of *guaBA* also decreased guanine pools in (p)ppGpp^0^ (Fig. 2D). Levels of ATP decreased in (p)ppGpp^0^ independent of deletion of *guaBA* (Fig. S2A & 3E), but levels of the other nucleotides differed only slightly between the strains (Fig. S2A-C). These results confirm a clear correlation between elevated guanine nucleotide levels during stationary phase starvation and decreased culturability.

### (p)ppGpp-deficient cells enter a division-incompetent but viable state during starvation

To further investigate the cause and mechanism of the decreased culturability of the (p)ppGpp^0^ mutant, we used different viability assays based on membrane integrity or metabolic activity. Despite the large decrease in CFUs during the stationary phase (Fig. 1B & S1A), only a small percentage of (p)ppGpp^0^ mutant cells were stained with the membrane-impermeable dye propidium iodide, and there was no significant difference between wildtype and the (p)ppGpp^0^ mutant (Fig. 3A). Analysis of the metabolic activity of late stationary phase cells by RedoxSensor™ Green staining, which monitors bacterial reductase activity, revealed slightly reduced metabolic activity in the (p)ppGpp^0^ mutant (Fig. 3B). However, the decrease in metabolic activity was less than expected from the difference in CFU counts between wildtype and (p)ppGpp^0^ during the late stationary phase (only approximately 10% culturable (p)ppGpp^0^ mutant). Resazurin conversion (Fig. 3C), an indicator of cellular redox potential, also revealed decreased respiratory activity in the (p)ppGpp^0^ mutant compared to wildtype and the *guaBA*-negative strain. To further confirm the alterations in the redox state of the mutant cells, we directly measured the NADH/NAD^+^ ratio (Fig. 3D) and ATP level (Fig. 3E). Both parameters were significantly decreased in the (p)ppGpp^0^ mutant. The effect was less prominent in the *guaBA* mutant background, indicating that the change in redox state is mainly a consequence of changes in the GTP pool.

**Fig. 3.**
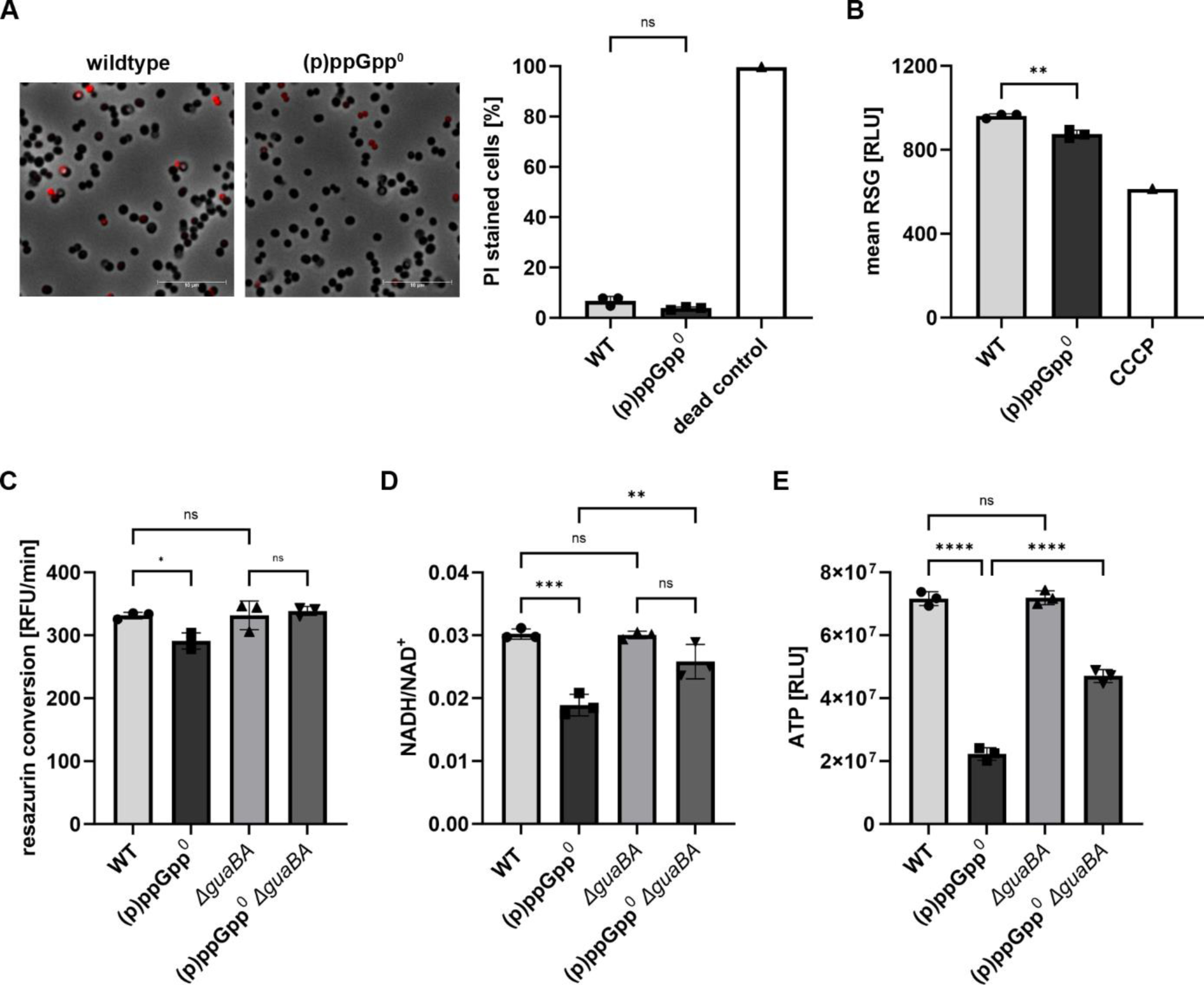
The (p)ppGpp^0^ mutant maintains membrane integrity but displays decreased metabolic activity. Strains were grown to late stationary phase (24h) in CDM. (A) Representative fluorescence microscopy images of propidium iodide (PI)-stained HG001 wildtype (WT) and (p)ppGpp^0^ are shown. The percentage of propidium iodide stained cells was measured by flow cytometry. For dead control bacteria were permeabilized with 70% EtOH. Data shown are mean ± SD (n=3 biological replicates, n=1 for dead control). Statistical significance was determined by one-way analysis of variance (ANOVA) with Tukey’s post-test (ns >0.05). Scale bar: 10µm (B) Bacteria were stained with RedoxSensor™ Green reagent and analysed by flow cytometry to monitor metabolic activity. Data shown are mean of RSG fluorescence ± SD (n=3 biological replicates, n=1 for CCCP control). Statistical significance was determined by one-way analysis of variance (ANOVA) with Tukey’s post-test (**p-value <0.01). (C) Redox potential was accessed by resazurin conversion over time in HG001 wildtype, (p)ppGpp^0^, *ΔguaBA* and (p)ppGpp^0^ *ΔguaBA.* Data shown are mean ± SD (n=3 biological replicates). Statistical significance was determined by one-way analysis of variance (ANOVA) with Tukey’s post-test (*p-value <0.05, ns >0.05). (D) NADH/NAD^+^ ratio and (E) relative ATP levels were determined. Data shown are mean of ± SD (n=3 biological replicates). Statistical significance was determined by one-way analysis of variance (ANOVA) with Tukey’s post-test (****p-value <0.0001, ***p-value <0.001, **p-value <0.01, ns >0.05).

The discrepancy between the decreased CFU counts and low propidium iodide staining, together with the decreased metabolic activity in the late stationary phase, suggest that the (p)ppGpp^0^ cells did not immediately lose viability. It is more likely that the cells entered a dormant state, slowing their metabolic activity and halting cell division, resulting in the observed large decrease in CFU.

### The division-incompetent state of (p)ppGpp^0^ cells is not the result of increased translation during the stationary phase

(p)ppGpp is known to efficiently inhibit protein synthesis ^9,26^ and thereby may prevent accumulation of protein aggregates ^27^. We measured protein synthesis at different time points during growth by incorporating the puromycin analogue O-propargyl-puromycin (OPP) into newly synthesized proteins. As expected, in the (p)ppGpp^0^ mutant, translation was significantly greater in the late exponential and early stationary growth phases (7 h and 9 h) (Fig. 4A). However, in the late stationary growth phase (24 h), complete shutdown of protein synthesis was observed, indicating that this occurred independently of (p)ppGpp. Delayed or absent translation inhibition can lead to formation of protein aggregates ^27^. However, only a slight increase in protein aggregates was observed in the (p)ppGpp^0^ mutant, as assessed by densitometric quantification of Coomassie-stained protein aggregates isolated from wildtype and (p)ppGpp^0^ late stationary phase cultures (Fig. S3A&B).

**Fig. 4.**
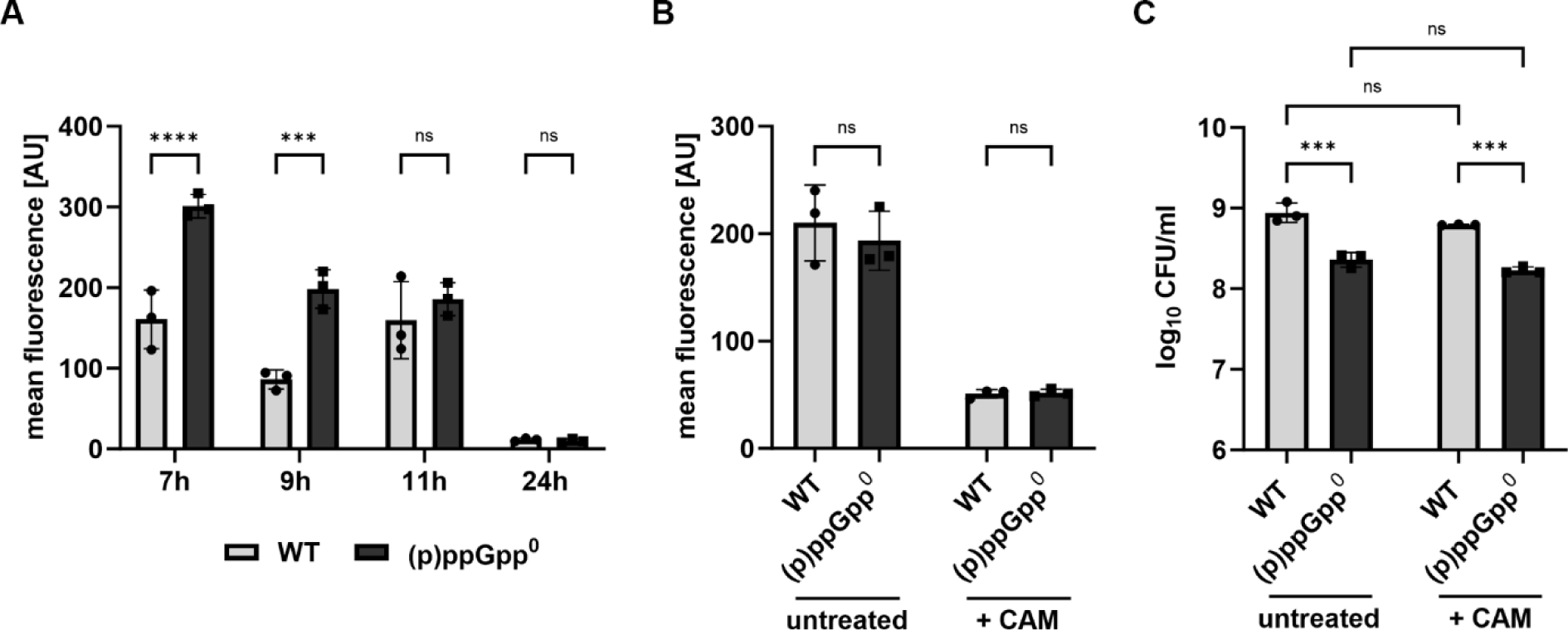
Increased translation of (p)ppGpp^0^ does not contribute to entrance into dormant state. (A) Protein synthesis was monitored at different timepoints during growth in wildtype (WT) and (p)ppGpp^0^ cells by labelling with the puromycin analogue OPP and subsequent flow cytometry analysis. Data shown are mean of ± SD (n=3 biological replicates). Statistical significance was determined by two-way analysis of variance (ANOVA) with Šidák’s post-test (****p-value <0.0001, ***p-value <0.001, ns >0.05). (B) Wildtype and (p)ppGpp^0^ cells grown to transition phase were treated with 100x MIC chloramphenicol (+ CAM), and protein synthesis by OPP incorporation and (C) culturability were assessed by CFU/ml enumeration. Data shown are mean of ± SD (n=3 biological replicates). Statistical significance was determined by two-way analysis of variance (ANOVA) with Šidák’s post-test (on log_10_ transformed data for (C), ***p-value <0.001, ns >0.05).

To further analyse whether increased translation in (p)ppGpp^0^ might contribute to entry into a division-incompetent state, we inhibited translation with 100x the MIC of the bacteriostatic antibiotic chloramphenicol. Chloramphenicol treatment almost completely inhibited protein synthesis in both wildtype and (p)ppGpp^0^ cells (Fig. 4B). However, chloramphenicol-induced inhibition of translation did not increase the culturability of (p)ppGpp^0^ cells (Fig. 4C).

Thus, translation inhibition appears to be delayed in the (p)ppGpp^0^ mutant, leading to slightly increased protein aggregation during the stationary phase. However, this delay in translation inhibition does not seem to be the cause of the nonculturable state of (p)ppGpp^0^ cells during stationary-phase starvation.

### Division-incompetent cell state of (p)ppGpp^0^ cells is not the result of oxidative stress during the stationary phase

In *S. aureus, (*p)ppGpp was shown to contribute to tolerance to oxidative stress ^28^, suggesting that elevated oxidative stress is a potential cause of the “dormant” state of (p)ppGpp^0^. To test this hypothesis, we analysed culturability under anaerobic and oxygen-limited, static (biofilm) conditions. Under these conditions, the culturability of the (p)ppGpp^0^ mutant was not compromised (Fig. 5A). However, treatment of aerobically grown cultures with the ROS scavenger N-acetyl cysteine (NAC) did not have any impact on culturability in the late stationary phase (Fig. 5B). Furthermore, no notable difference in endogenous ROS formation was detected between wildtype and the (p)ppGpp^0^ mutant during the stationary growth phase (Fig. 5C). Thus, (p)ppGpp-deficient cells were impaired only in culturability under aerobic conditions. However, this phenotype could not be linked to elevated ROS levels. Thus, we investigated the contributions of metabolic cues such as aerobic respiration and proton motive force (PMF).

**Fig. 5.**
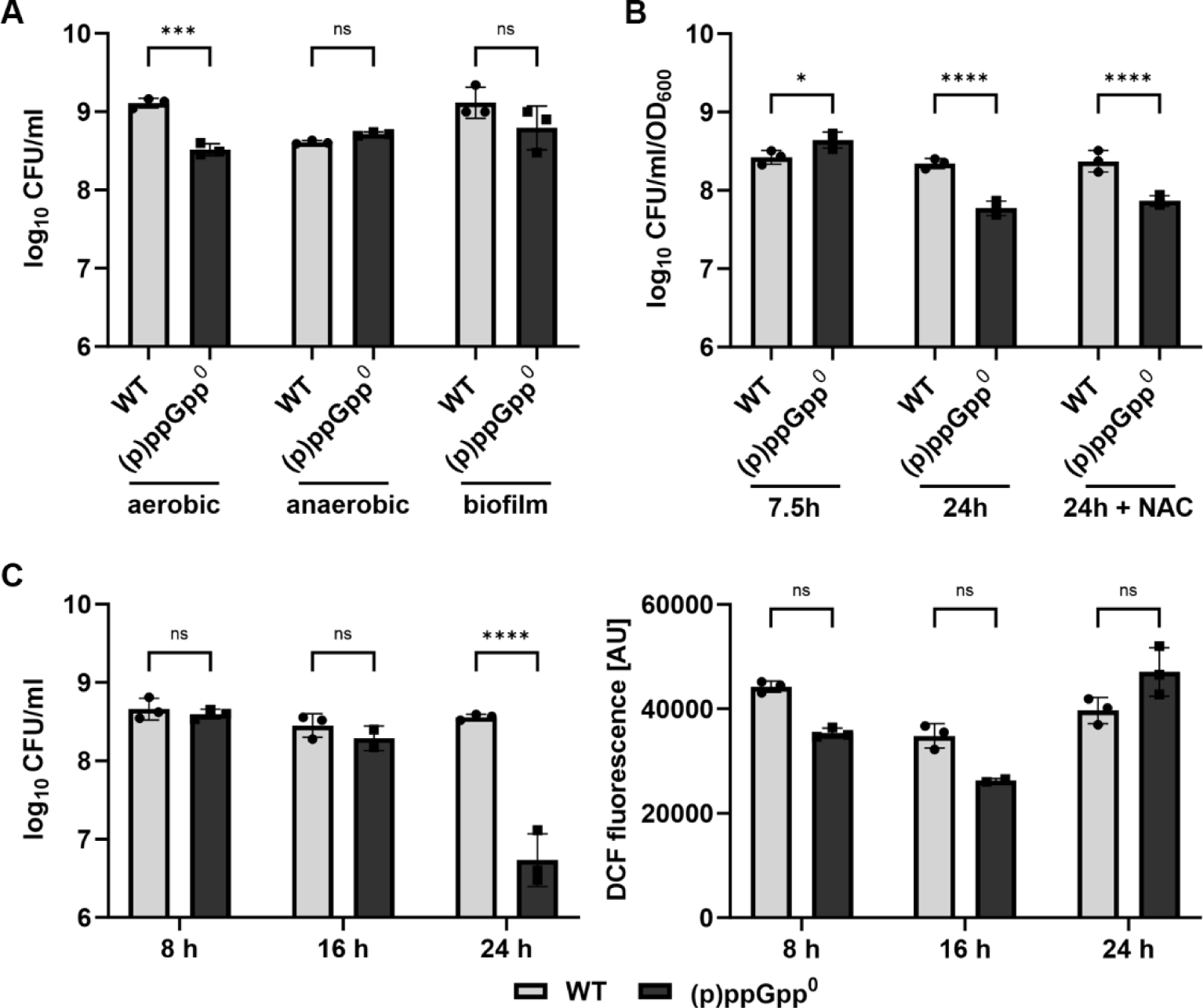
Oxidative stress in (p)ppGpp^0^ does not contribute to entrance into dormant state. (A) HG001 wildtype (WT) and (p)ppGpp^0^ strain were grown under aerobic, anaerobic or oxygen-limited, static (biofilm) conditions for 24h and culturability was accessed by CFU/ml enumeration. Data shown are mean ± SD (n=3 biological replicates). Statistical significance was determined by two-way analysis of variance (ANOVA) with Šidák’s post-test on log10 transformed data (***p-value <0.001, ns >0.05). (B) Wildtype and (p)ppGpp^0^ cells were treated with 12.5 mM N-acetylcysteine (+ NAC) during transition to stationary phase (7.5h) and CFU/ml counts normalized to OD_600_ were determined before and after treatment. Data shown are mean of ± SD (n=3 biological replicates). Statistical significance was determined by two-way analysis of variance (ANOVA) with Šidák’s post-test (****p-value <0.0001, * <0.05). (C) Monitoring of culturability, as determined by CFU/ml enumeration, and intracellular ROS levels, as measured by using the DCFH2-DA dye, which is oxidized to DCF by ROS, from early to late stationary phase. Data are shown as mean ± SD (n=3 biological replicates). Statistical significance was determined by two-way analysis of variance (ANOVA) with Šidák’s post-test (****p-value <0.0001, ns >0.05).

### Membrane function and architecture are impaired in (p)ppGpp^0^ cells

Changes in membrane potential can impact cell vitality by perturbing ATP synthesis and ion transport. We measured membrane potential by staining with the carbocyanine dye DiOC2(3). During the exponential growth phase, there was no difference in membrane potential between wildtype, (p)ppGpp^0^ and *guaBA-*negative cells (Fig. 6A). However, during the late stationary phase, the membrane potential of the (p)ppGpp^0^ mutant was notably lower than that of wildtype. In contrast, there was no significant difference in the level of the *guaBA*-negative strain, indicating that high levels of GTP disturb the membrane potential. The PMF is composed of the electrical potential ΔΨ and the transmembrane proton gradient ΔpH. The intracellular pH was similar between wildtype and the (p)ppGpp^0^ mutant (Fig. 6B), indicating that membrane potential ΔΨ mainly reduced in cells with high GTP levels. Analysis of the membrane and cell wall architecture by costaining with the dye FM4-64X and the BODIPY-vancomycin conjugate revealed irregular staining of the (p)ppGpp^0^ cell membrane during starvation in the stationary phase, which was not observed in wildtype cells (Fig. 6C). In contrast, no abnormalities were detected in the cell wall of (p)ppGpp^0^ cells during stationary phase starvation compared to wildtype cells (Fig. 6C). These irregularities in the cell membrane of the (p)ppGpp^0^ strain, together with increased membrane fluidity, as measured by DPH fluorescence polarization (Fig. 6D), are indicative of membrane patches with increased fluidity, also known as regions of increased fluidity (RIFs).

**Fig. 6.**
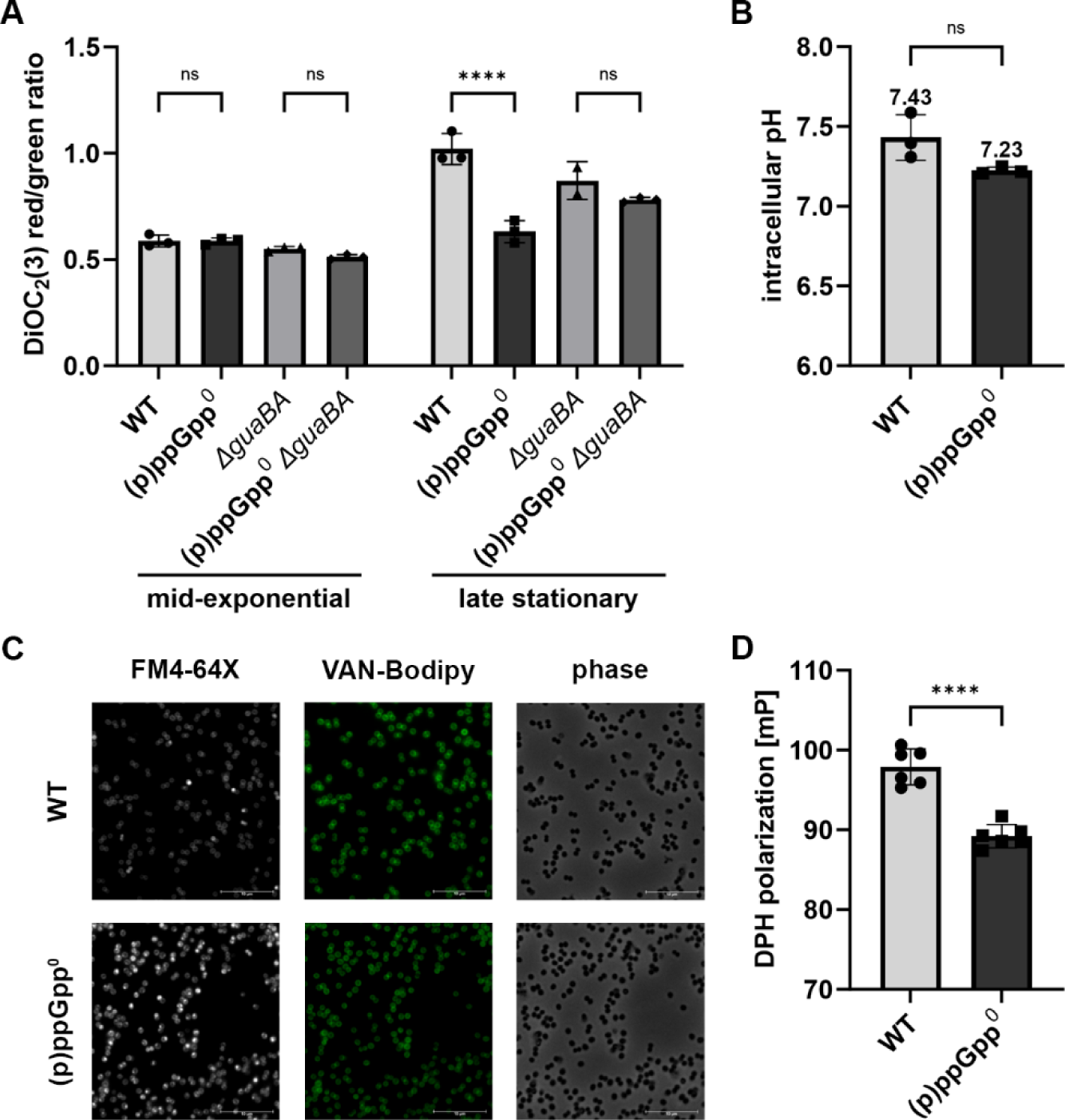
Membrane function and architecture is altered in the (p)ppGpp^0^ mutant. (A) Membrane potential was measured during mid-exponential and late stationary phase by staining with the carbocyanine dye DiOC_2_(3). Data shown are mean of ± SD (n=3 biological replicates). Statistical significance was determined by two-way analysis of variance (ANOVA) with Šidák’s post-test (****p-value <0.0001, ns >0.05) (B) Intracellular pH was determined by pHRodo Green AM staining of late-stationary phase bacteria. Data shown are mean ± SD (n=3 biological replicates). Statistical significance was determined by a two-tailed, unpaired t-test (ns > 0.05). (C) Representative microscopic images of FM4-64X (cell membrane) and BODIPY-vancomycin conjugate (cell wall) double-stained *S. aureus* from late stationary phase (24h). Scale bar: 10µm (D) Membrane fluidity was measured by DPH fluorescence polarization in late stationary phase (24h). Data shown are mean ± SD (n=6 biological replicates). Statistical significance was determined by a two-tailed, unpaired t-test (****p-value <0.0001).

Taken together, the results show that in the stationary phase, high levels of GTP, as found in the (p)ppGpp^0^ mutant, led to decreased membrane potential, changes in cell membrane architecture and increased membrane fluidity. Thus, the decreased culturability of the (p)ppGpp^0^ mutant is likely due to loss of membrane potential.

### Impact of a lack of (p)ppGpp biosynthesis and GTP imbalance on the transcriptome during stationary phase starvation

To better understand how changes in GTP levels might alter membrane potential, we analysed changes in gene expression in wildtype (GTP low), (p)ppGpp^0^ mutant (GTP high) and *ΔguaAB* mutant (GTP low) cells after 24 h of growth (Supplementary Tables S1 und S2). Genes with at least a log2-fold change of ≥ 1 and ≤ -1 and a p value ≤ 0.001 were defined as differentially regulated. In total, 235 and 288 genes were significantly upregulated or downregulated, respectively, in the (p)ppGpp^0^ mutant compared to wildtype. Such (p)ppGpp-dependent and GTP-independent differences were much less prominent in the *guaBA* background (only 50 and 65 genes up- or downregulated, respectively). To assess (p)ppGpp-independent and GTP-dependent changes in gene expression, the (p)ppGpp^0^ *ΔguaBA* mutant was compared to the (p)ppGpp^0^ mutant, and 514 and 628 genes were found to be significantly up- or downregulated, demonstrating that under our experimental conditions, the majority of transcriptomic changes are mediated by differences in GTP levels, as opposed to being direct effects of (p)ppGpp.

Many of the previously identified stringent response genes identified after challenge with amino acid limitation ^12^ or transcriptional induction of (p)ppGpp synthetases ^22^ were differentially regulated during stationary phase starvation (see Fig. S4 and S1C). Most CodY target genes were downregulated in the (p)ppGpp^0^ mutant [e.g., *rsaD* (-185.8-fold), the *cap* operon (-5,2-fold for *capA*) and *aur* (-65,7-fold)]. Likewise, genes involved in amino acid synthesis and transport, such as *brnQ2* (-6.1-fold), *gltBD* (-14.8-fold and -12.2-fold), *dapABDL* (-12.5-fold for *dapA*), the *opp-3ABCDEF* operon (-20.1-fold for *opp-3B*) and *ilvD* (-7.2-fold), were also downregulated. Furthermore, several genes of the SigB regulon were downregulated in (p)ppGpp^0^. For instance, *krtA*, which encodes a potassium transporter, was downregulated 2.1-fold. Similarly, *clpL*, which is involved in protein metabolism, was downregulated 6.1-fold. The *mnh* operon, which is hypothesized to encode a redox-energized complex involved in PMF maintenance ^29^, was also downregulated 5.1-fold. Furthermore, *groEL* (2.4-fold) and *groES* (4.9-fold), which are part of the CtsR/HrcA regulon, were found to be upregulated, indicative of proteotoxic stress. Both protein complexes were upregulated during proteotoxic stress, which leads to protein misfolding and aggregation ^30^. These findings are consistent with the slightly elevated protein aggregates detected in the (p)ppGpp^0^ mutant during stationary phase starvation (Fig. S4F). Interestingly, phage-encoded genes were upregulated under GTP-high conditions, indicating that phages φ11, φ12 and φ13 are induced in (p)ppGpp^0^ during the stationary phase. These observations are in agreement with previous studies that found the ΦSa3 phage to be downregulated under guanine-limited conditions ^31^. Notably, the transcriptome further revealed strong downregulation of TCA cycle (*citB*-3.1-fold, *sucA*-2.3-fold and *sucB*-2.6-fold) and aerobic respiration (*qoxABCD* -7.6-fold for *qoxA*, *ctaA* -3.4-fold and ATP synthase e.g. *atpD* -2,3-fold) genes (Fig. 7), which was GTP dependent, as both the TCA cycle and respiration were mostly unaffected in the *guaBA* negative background (Fig. 7). Downregulation of the TCA cycle and respiration may account for the reduced membrane potential observed during stationary phase starvation in the (p)ppGpp^0^ mutant (Fig. 6A).

**Fig. 7.**
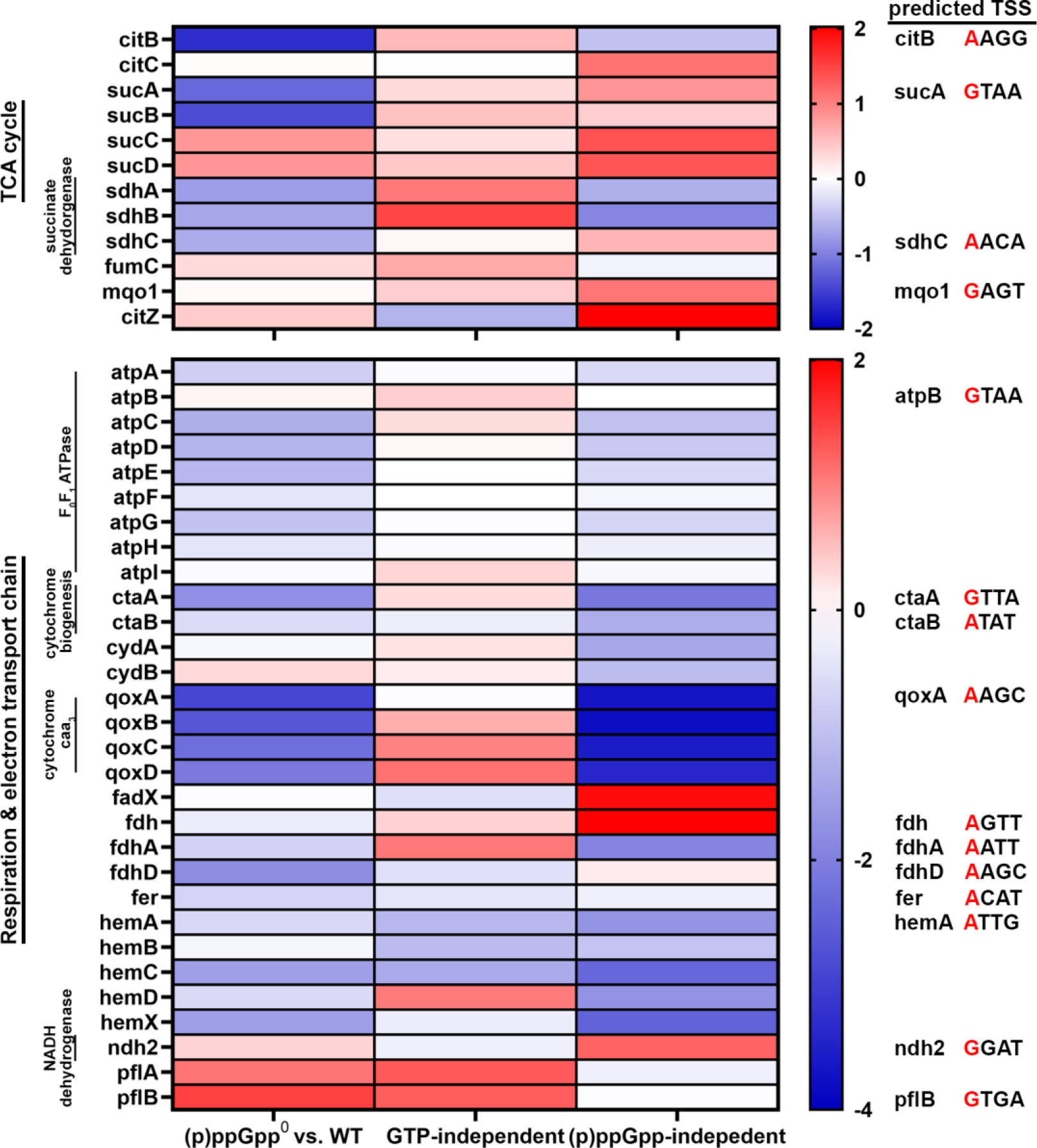
Transcriptomic changes in TCA cycle and respiration in dormant (p)ppGpp^0^ cells. Heatmaps displaying the RNA-seq fold change of selected genes involved in the TCA cycle and electron transport chain, and related to respiration (SEED annotation). Log_2_ fold changes in relative transcript abundances are colour-coded with red and blue, indicating up- and downregulation, respectively. “GTP-independent” compares the expression levels of (p)ppGpp^0^ *ΔguaBA* vs. *ΔguaBA*, while “(p)ppGpp-independent” compares the expression levels of (p)ppGpp^0^ vs. (p)ppGpp^0^ *ΔguaBA*. The predicted TSS position between nucleotide +1 and +4 is depicted, while the TSS +1 is indicated in red colour. The RNA-seq data shown are from three individual biological replicates (n=3). Full RNA-seq data are available in Data S1 and S2 in the supplementary material.

### The sensitivity of *qoxABCD* expression to GTP determines the activity of the electron transport chain and PMF

Since the transcriptome analysis of the (p)ppGpp^0^ mutant revealed not only downregulation of the TCA cycle but also strongly decreased expression of the terminal oxidase gene *qoxABCD*, a component of the electron transport chain (ETC), we hypothesized that these transcriptional changes lead to decreased PMF.

We verified the RNAseq data by RT‒qPCR and confirmed decreased *qoxA* expression in (p)ppGpp^0^ but increased expression in the *guaBA*-negative strain compared to wildtype (Fig. 8B). Expression of rRNA is controlled by the concentration of the transcription-initiating nucleotide (iNTP) within the promoter ^11^. Analysis of potential transcriptional start sites upstream of *qoxABCD* predict an ATP as the transcription-initiating nucleotide (based on data from ^32^)(Fig. 7). Thus, we mutated the TSS+1 to a GTP in wildtype and (p)ppGpp^0^ (named WT PqoxABCD_mut_ and (p)ppGpp^0^ PqoxABCD_mut_) (Fig. 8A). This mutated TSS+1 increased *qoxA* expression in the late stationary phase in (p)ppGpp^0^, indicating that this promoter is nucleotide sensitive (Fig. 8B). Alteration of the iATP to iGTP also resulted in greater CFU counts than those of (p)ppGpp^0^ (Fig. 8C). However, culturability was still lower compared to wildtype, showing that additional factors are likely to contribute to entry into dormancy.

**Fig. 8.**
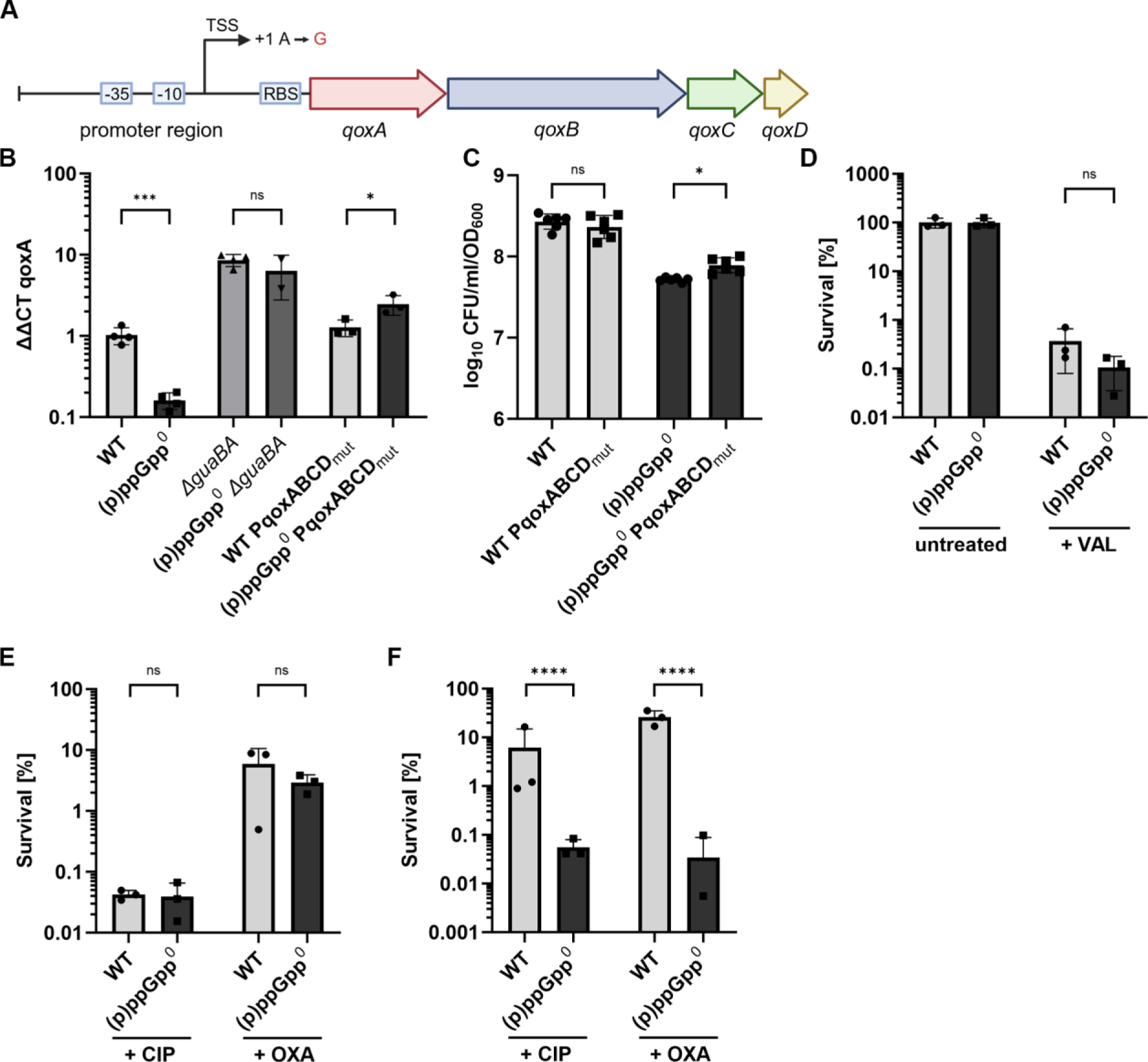
Decrease in ETC activity contributes to entrance into dormant state and decreased antibiotic tolerance (A) *qoxABCD* gene operon with promoter region. The predicted initiation nucleotide (TSS +1) iATP was mutated to iGTP. (B) Bacterial cells were harvested during the late stationary growth phase and total RNA was isolated. *qoxA* transcript levels were determined by RT-qPCR and normalized to *gyrB* expression by ΔΔCt method. Data are shown as mean ± SD (n=3 biological replicates). Statistical significance was determined by two-tailed unpaired t-tests (***p-value <0.001, *p-value <0.05, ns >0.05). (C) Culturability during late stationary phase (24h) was determined by CFU enumeration and normalized to OD_600_. Data shown are mean of ± SD (n=6 biological replicates from two individual experiments). Statistical significance was determined by a one-way analysis of variance (ANOVA) with Tukey’s post-test (*p-value <0.05). (D) Wildtype (WT) and (p)ppGpp^0^ cells were treated with a sub-inhibitory concentration of valinomycin (20 µM) during the late stationary growth phase (24h) and viability was assessed by CFU enumeration after 24h of treatment. Survival was calculated in comparison to untreated samples. Data shown are mean of ± SD (n=3 biological replicates). Statistical significance was determined by a two-way analysis of variance (ANOVA) with Šidák’s post-test (ns >0.05, based). (E) Mid-exponential phase or stationary phase (F) cells were treated with 100x MIC ciprofloxacin (+ CIP) or oxacillin (+ OXA) and survival was calculated after 3h (E) or 24h (F) of treatment in comparison to untreated cells. Data shown are mean of ± SD (n=3 biological replicates). Statistical significance was determined by a two-way analysis of variance (ANOVA) with Šidák’s post-test (****p-value <0.0001, ns >0.05).

To further demonstrate the importance of the PMF for stationary phase survival, we treated wildtype and (p)ppGpp^0^ cultures with subinhibitory concentrations of valinomycin during the late stationary phase and evaluated survival after 24 h. Valinomycin is an ionophore that disrupts membrane potential by selectively translocating potassium ions across the membrane. Upon valinomycin treatment, wildtype and (p)ppGpp^0^ showed similar reductions in CFU counts compared to untreated cultures (Fig. 8D). Thus, disruption of membrane potential indeed compromises culturability and survival.

Together, these findings illustrate the significance of preserving PMF, particularly ΔΨ, throughout stationary phase starvation to sustain culturability.

### Preservation of PMF during starvation is crucial for antibiotic tolerance

It has been demonstrated that maintenance of PMF is essential for the survival of starvation-induced tolerant bacteria ^33,34^. We sought to determine whether the decreased membrane potential in the (p)ppGpp^0^ mutant would also result in a reduction of starvation-induced antibiotic tolerance. Therefore, we cultured wildtype and (p)ppGpp^0^ cells to exponential or stationary growth phase and exposed them to either 100x MIC oxacillin or ciprofloxacin. No significant difference in antibiotic survival between wildtype and (p)ppGpp^0^ cells was observed when treated during exponential growth (Fig. 8E). However, wildtype cells were barely killed by the antibiotics during the stationary phase, confirming the higher antibiotic tolerance of non-growing bacteria ^3^. However, both antibiotics efficiently decreased the survival of the (p)ppGpp^0^ mutant during the stationary phase (Fig. 8F).

## Discussion

Here, we show that bacterial culturability in the stationary phase is dependent on (p)ppGpp-driven GTP depletion. Uncontrolled, high levels of GTP in the (p)ppGpp^0^ mutant decrease membrane potential, at least in part, via transcriptional downregulation of nucleotide-sensitive genes of the TCA cycle and the respiratory chain. Maintenance of respiratory chain activity by (p)ppGpp/low GTP) is also responsible for the tolerance of stationary phase cells to bactericidal antibiotics (Fig. 9).

**Fig. 9.**
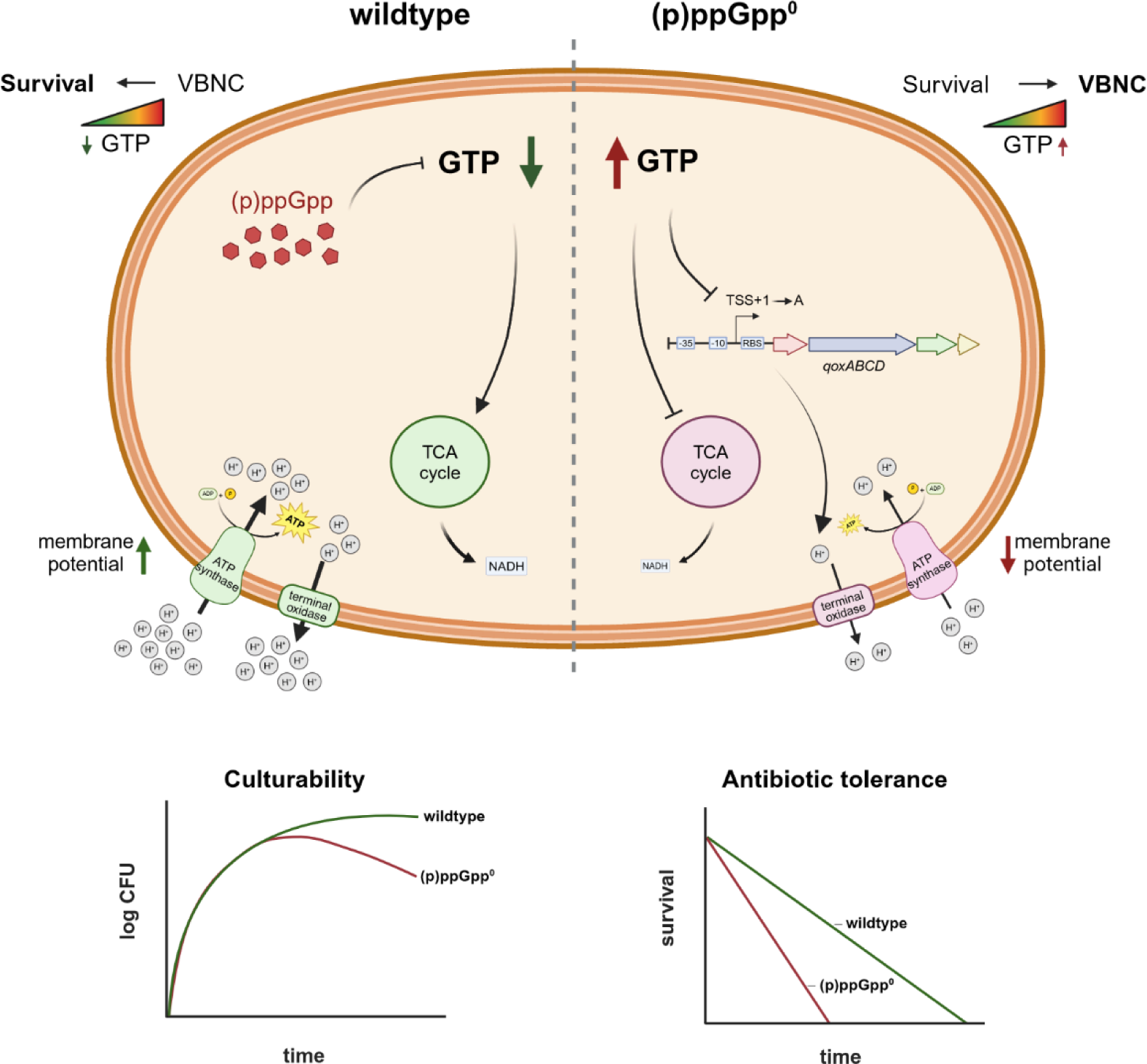
(p)ppGpp-dependent regulation of GTP homeostasis enables maintenance of culturability. During stationary phase starvation, (p)ppGpp synthesis contributes to regulation of intracellular GTP levels and thereby maintains membrane potential and ensures culturability and survival during antibiotic treatment. In a (p)ppGpp-deficient mutant, GTP homeostasis is disturbed leading to decreased expression of the TCA cycle and electron transport chain (ETC) genes including the terminal oxidase *qoxABCD*. This in turn leads to a decreased membrane potential which causes entrance into a division-incompetent but viable (VBNC) state, accompanied by reduced antibiotic tolerance.

### (p)ppGpp-dependent GTP homeostasis determines long-term bacterial culturability

The defect in culturability of the (p)ppGpp^0^ mutant is clearly linked to an uncontrolled increase in GTP levels upon entry into the stationary phase. Culturability correlated inversely with changes in the GTP level induced by deletion of *purR* or *guaAB* or by guanine feeding. Mutation of *guaAB* rescued the defect in the (p)ppGpp^0^ mutant. A high guanine concentration did not interfere with the survival of wildtype, indicating that (p)ppGpp can also control GTP levels provided by the salvage pathway. Direct (p)ppGpp-dependent alterations in translation or replication machinery ^9,15^ are likely to also be less prominent in a *guaAB*-negative background. Thus, we conclude that the level of GTP homeostasis is the main determinant of bacterial culturability in stationary phase cells.

Control of guanine metabolism by (p)ppGpp is conserved in several Firmicutes species, such as *B. subtilis*, ^13^, *E. faecalis* ^35^ and *S. aureus* ^14^. Recently, reduced long-term survival of the *S. aureus* USA300 (p)ppGpp^0^ mutant ^18^ was linked to disrupted GTP homeostasis since suppressor mutants with alterations in *gmk* were selected during long-term culture. In *B. subtilis*, low GTP levels reduce the growth rate but promote survival during amino acid starvation ^13,36^ and cell viability during fatty acid starvation ^37^. Elevated GTP levels in a (p)ppGpp^0^ mutant of *B. subtilis* promoted the bactericidal effect of otherwise bacteriostatic antibiotics ^38^.

### Disturbance of GTP homeostasis during nutrient stress leads to induction of a VBNC state

Previously, it was proposed that (p)ppGpp deletion in *B. subtilis* results in “death-by-GTP” and that this cell death might be due to unrestricted translation ^13^. However, in *S. aureus*, uncontrolled translation could not be linked to diminished bacterial survival since translation inhibitors did not restore culturability of the (p)ppGpp^0^ mutant. Furthermore, our results do not support the hypothesis that high GTP levels trigger a classical “cell death” programme. The (p)ppGpp^0^ mutant of *S. aureus* was nonculturable but negative for propidium iodide uptake and maintained basal metabolic activity and transcription, albeit with significantly reduced ATP levels. This is indicative of the previously defined state of VBNC (viable but nonculturable) or a deep dormant-like state ^39^. It was proposed that ATP depletion is involved mainly in formation of VBNCs ^27,40^, in which intracellular energy (ATP) is the main factor maintaining the culturability of bacteria on solid media. In some gram-negative bacteria, (p)ppGpp was shown to prevent VBNC state formation in response to certain stress conditions ^41,42^. In (p)ppGpp^0^ mutants of *B. subtilis*, fatty acid starvation leads to membrane rupture and, consequently, cell death ^37^. The authors proposed a model whereby during fatty acid starvation, a (p)ppGpp-dependent decrease in GTP levels halts growth, preventing collapse of membrane potential and loss of membrane integrity. However, nonculturable (p)ppGpp^0^ *B. subtilis* mutants died, as indicated by propidium iodide staining ^37,38^. Thus, there might be major differences between GTP-dependent cell death pathways in *S. aureus* and *B. subtilis*.

### How do elevated GTP levels lead to VBNC-state formation?

Our RNAseq analysis revealed that TCA cycle genes and genes coding for major components of the electron transport chain, including the terminal oxidase (qoxABCD) and the F0F1 ATP synthase, were significantly downregulated in the (p)ppGpp^0^ mutant during the stationary phase. Accordingly, ATP levels (Fig. 3E), the NADH/NAD+ ratio and membrane potential (Fig. 6A) were decreased in the (p)ppGpp^0^ mutant. In Firmicutes, (p)ppGpp-dependent transcriptional changes are regulated mainly by the GTP-dependent transcriptional repressor CodY ^20^ or by the availability of the NTP substrate for transcription initiation ^11,14,43^. We could exclude any influence of CodY (Fig. 2B) on induction of the VBNC state. We mutated the predicted initiation of NTP transcription in the promoter region of *qoxABCD*, the most prominently downregulated component of the electron transport chain. By using this approach, we were able to restore wildtype levels of *qoxA* expression in the (p)ppGpp^0^ mutant (Fig. 8B). Furthermore, the culturability of the (p)ppGpp^0^ mutant was partially restored by this single nucleotide exchange. Expression of other genes involved in respiration and generation of membrane potential was downregulated in the (p)ppGpp^0^ mutant, which was also initiated with iATP (Fig. 7); these genes may further contribute to the observed phenotype.

### PMF maintenance during starvation is crucial for antibiotic tolerance

We showed that a (p)ppGpp^0^ mutant grown to the stationary phase lost antibiotic tolerance to the β-lactam oxacillin and the fluoroquinolone antibiotic ciprofloxacin. Nongrowing wildtype bacteria are tolerant to these antibiotics, as shown previously ^3^). However, Conlon et al. did not find evidence that (p)ppGpp is involved in antibiotic persistence or tolerance. The authors proposed that ATP depletion is important for the observed tolerance of stationary phase cells

### 3. This seems to contrast our findings that (p)ppGpp-dependent maintenance of low GTP/high

ATP levels protected the cells. This discrepancy might be explained by the experimental design. The growth medium used by Conlon et al. ^3^ likely does not induce a stringent response and/or might contain low amounts of guanine. Thus, under these conditions, the intracellular GTP pool might not be elevated in the (p)ppGpp^0^ mutant. In agreement with our results, previous studies have shown that PMF is essential for starvation-induced tolerance in both gram-negative and gram-positive bacteria, including *S. aureus* ^33,34,44^. Hence, these findings support the idea of applying PMF inhibitors to combat antibiotic-tolerant bacteria, including *S. aureus* ^45,46^.

## Materials and Methods

### Growth conditions and CFU determination

The strains and plasmids used in this study are listed in the Supplementary Information, Table S1 and S2. If not indicated elsewhere, *S. aureus* strains were grown in LB-Miller, tryptic soy broth (TSB) or chemically defined medium (CDM) ^47^ at 37 °C and 200 rpm.

For CFU experiments, bacteria from an overnight culture grown in LB-Miller were diluted to an initial optical density at 600 nm (OD_600_) of 0.05 in fresh medium without antibiotics and grown with shaking (200 rpm) at 37 °C to the desired growth phase. LB-Miller medium was used overnight culture since the (p)ppGpp^0^ mutant in this medium showed no defect in survival. For oxygen-limited growth (biofilm conditions), bacteria were inoculated into 1 ml of fresh medium in a 24-well plate and statically grown at 37 °C. For anaerobic growth conditions, 14 ml of fresh medium was inoculated into glass tubes, which were closed with a rubber stopper to minimize the oxygen supply and subsequently grown at 37 °C and 200 rpm. For certain CFU experiments, 400 µg/ml chloramphenicol (100x MIC of HG001 wildtype and (p)ppGpp^0^), 12.5 mM N-acetylcysteine or 20 µM valinomycin was added at the indicated timepoints during growth. For CFU/ml determination, a 10-+fold dilution series was prepared in phosphate-buffered saline (PBS), 10 µl of each dilution was spotted onto TSB agar, and CFU counts were determined after 16–18 h of incubation at 37 °C.

### Generation of *S. aureus* mutant strains

#### Generation of SH1000 (p)ppGpp^0^

The mutagenesis strategy described by Geiger et al. was used for generation of the SH1000 (p)ppGpp^0^ mutant (SH1000-229-230-263) ^20^. Lysates were prepared from RN4220 strains containing the mutagenesis vectors pCG229, pCG230 and pCG263. After plasmid transduction, mutagenesis was performed as previously described ^48^. The mutations were verified by PCR and sequencing using the oligonucleotides listed in Table S3.

#### Transduction of purR mutants

Φ11 lysates were generated from the transposon mutant NE1237 (purR) of the NARSA transposon library ^49^ to transduce *S. aureus* HG001 wildtype and (p)ppGpp^0^ cells. The mutations were verified via PCR using the oligonucleotides listed in Table S3.

#### Mutagenesis strategy for WT-PqoxABCD_mut_ and (p)ppGpp^0^-PqoxABCD_mut_

The plasmid pCG919mut was constructed for base substitution at the position +1 of the transcriptional start site of the P*_qoxABCD_* promoter. The complete promoter region with approximately 1000 bp of the left and right flanking regions was amplified in two PCRs using the primers pCG919gibfor and pCG919-2mutfor and pCG919-2mutrev and pCG919gibrev. The PCR products were subsequently cloned and inserted into the Bam HI-digested pIMAY-Z vector via Gibson cloning and transformed into *E. coli* IM08B. After verification of the base substitution (A → G) by sequencing, the plasmid was transformed into *S. aureus* HG001 wildtype and (p)ppGpp^0^ cells by electroporation. The mutation was introduced into the genome by pIMAY mutagenesis ^50,51^ and verified by sequencing. All oligonucleotides used are listed in Supplementary Table S3.

### RNA isolation, qRT‒PCR and RNAseq

RNA isolation and qRT‒PCR were performed as described previously ^22^. Briefly, bacteria were pelleted, resuspended in 1 ml of TRIzol (Thermo Fisher Scientific) and lysed using zirconia/silica beads (0.1 mm diameter) and a high-speed homogenizer. RNA was isolated following the procedure recommended by the TRIzol manufacturer.

Relative quantification of *psmα, rsaD, rpsL* and *qoxA* transcripts by qRT‒PCR was performed with QuantiFast SYBR Green RT‒PCR Kit (Qiagen) using the Quantstudio3 system (Applied Biosystems). Briefly, 5 μg of total RNA was DNase-treated and diluted 1:10 for qRT‒PCR. Relative expression of transcripts was calculated with the ΔΔCT method and normalized to *gyrB* gene expression. The sequences of the oligonucleotides used for qRT‒PCR are listed in Table S3.

For RNA-seq analysis, RNA isolated from the aqueous phase was further purified with an Amp Tech ExpressArt RNA Reading Kit. For each sample, a total of 100 ng of RNA was subjected to rRNA depletion, followed by cDNA library construction via IlluminaTM Stranded Total RNA Prep Ligation with a Ribo Zero Plus Kit according to the manufacturer’s instructions. Libraries were sequenced as single reads (100 bp read length) using the NextSeq platform (Illumina) and NextSeq 500 Mid Output Kit v2.5. The sequences were demultiplexed with bcl2fastq (v2.19.0.316), quality checked with fastq (v0.20.1) and visualized with MultiQC (v1.7). Analysis of the RNAseq results was performed using CLC Genomic Workbench 23.0.1 (Qiagen). Reads were mapped to the reference genome of HG001 (NZ_CP018205.1). Differential gene expression was performed comparing (p)ppGpp^0^ vs. wildtype, (p)ppGpp^0^ *ΔguaBA* vs. *ΔguaBA* (GTP independent) and (p)ppGpp^0^ vs. (p)ppGpp^0^ *ΔguaBA* ((p)ppGpp independent). Genes with at least log_2_ fold change of >1 or <-1 and a p-value <0.001 were defined as differentially regulated. Annotation of genes was performed according to the recent “Aureowiki” annotation of strain 8325 (http://aureowiki.med.unigreifswald.de/Main_Page) ^52^.

### Metabolite extraction and LC‒MS/MS quantification

For metabolite extraction, 3 ml of bacterial culture was harvested by vacuum filtration, washed twice with 0.6% NaCl solution, transferred directly into 5 ml of quenching solution (acetonitrile:methanol:water, 40%:40%:20%, (v/v/v)) and shock frozen in liquid nitrogen. Afterwards, 1 ml of the cell suspension was lysed using zirconia/silica beads (0.1 mm diameter) and a high-speed homogenizer. The lysate was centrifuged at -9 °C at 20 000 × g for 15 minutes, and the clear supernatant was stored at -80 °C for LC–MS/MS analysis. Relative concentrations of the nucleotides in the cell extracts were measured via isotope ratio LC– MS/MS as described previously ^53^. For evaluation, the 12C/13C ratios were normalized to the OD_600_ of the bacterial culture, and the relative abundances of each metabolite are shown.

### LIVE/DEAD and viability staining using microscopy and flow cytometry

Bacterial cultures were concentrated 10-fold and washed in 0.85% NaCl prior to staining for bacterial viability, as recommended by the manufacturer (Live/Dead BacLight^TM^ Bacterial Viability Kit). All cultures were further diluted 5-to 10-fold to an OD_600_ of 0.2-0.4 before staining with the Syto 9 and propidium iodide dye mixture (1:1 dye ratio). Fluorescence microscopy was performed with a Leica DM5500B fluorescence microscope using the following filter sets (excitation and emission filters): Syto 9: BP470 40-nm and BP525 50-nm; propidium iodide: BP535 50-nm and BP610 75-nm. Images were captured using a Leica DFC360FX monochrome camera with the following exposure times: 40 ms for Syto 9, 150 ms for propidium iodide and 6 ms for phase contrast channels. Images were viewed with in-built Leica ASF software. Alternatively, the percentage of propidium iodide-stained cells was analysed via flow cytometry using a FACSCalibur flow cytometer (Becton Dickinson).

### RedoxSensor Green assay

To monitor bacterial reductase activity as an indicator of electron transport chain function, cells from the stationary phase were stained with RedoxSensor™ Green reagent (BacLight™ RedoxSensor™ Green Vitality Kit) and analysed by flow cytometry according to the manufacturer’s instructions.

### AlamarBlue measurement

To measure cell viability via resazurin conversion, a stationary phase culture was tenfold diluted in PBS and mixed according to the manufacturer’s instructions with alamarBlue™ reagent at a 1:10 ratio. After incubating at 37 °C, resofurin fluorescence (excitation: 540 nm, emission: 600 nm) was measured every 5 min. The increase in resofurin fluorescence over time was analysed via simple linear regression in GraphPad Prism 10.

### ATP and NADH/NAD^+^ ratio determination

ATP levels and the NADH/NAD^+^ ratio were determined using the BacTiter-Glo™ Microbial cell Viability assay or the NAD/NADH-Glo™ assay (Promega) following the manufacturer’s instructions. Luminescence was measured in a white 96-well plate using the Tecan Spark luminescence module.

### Click-it OPP protein synthesis labelling

Protein synthesis was quantified with a Click-it™ Plus OPP Alexa Fluor™ 488 protein synthesis labelling kit according to the manufacturer’s instructions. Briefly, growing bacterial cells were incubated with 20 µM O-propargyl puromycin (OPP) for 30 min. Afterwards, the cells were harvested by centrifugation, fixed with formaldehyde and permeabilized with 70% ethanol (v:v) and lysostaphin. Then, OPP was conjugated to Alexa Fluor™ 488 using click chemistry according to the manufacturer’s protocol. The stained cells were analysed using flow cytometry (Beckton Dickinson FACS Calibur).

### Detection and quantification of protein aggregation

Protein aggregates within cells were isolated according to a described protocol ^54,55^. Briefly, protein aggregates were isolated from 5 ml of culture. First, bacterial cells were harvested by centrifugation and washed twice with PBS. Then, the cells were resuspended in buffer A (50 mM Tris, 150 mM NaCl, pH 8) and lysed using zirconia/silica beads (0.1 mm diameter) and a high-speed homogenizer at 6 m/s three times for 30 seconds each. The crude extract was centrifuged at 18 000 × *g* for 30 minutes, after which the pellet was washed twice with buffer A containing 0.5% Triton X-100. The protein aggregates were then solubilized in rehydration buffer (7 M urea, 2 M thiourea, 4% (wt/vol) CHAPS, 100 mM DDT) and loaded on a 12% SDS‒ PAGE gel. Afterwards, the SDS‒PAGE mixture was stained with InstantBlue™ Coomassie blue-based staining solution (expedeon). Protein aggregates were densitometrically quantified from Coomassie blue-stained gels using the GelAnalyzer plugin of ImageJ software.

### ROS measurement

Endogenous ROS levels were measured using 2′,7′-dichlorodihydrofluorescein diacetate (DCFH2-DA) dye based on a previous protocol ^56^. DCFH2-DA is deacetylated by alkaline hydrolysis to generate DCFH2, which is oxidized by ROS to the fluorescent dye 2′,7′-dichlorofluorescein (DCF). Briefly, *S. aureus* HG001 wildtype and (p)ppGpp^0^ cells were cultivated in CDM for the indicated durations and harvested at an OD600 equivalent of 3 × 10^8^ cells by centrifugation. The cell pellets were incubated with DCFH2 for 40 min as previously described [75]. Relative DCF fluorescence was measured using a plate reader (Tecan Spark), with an excitation wavelength of 480 nm with a bandwidth of 20 nm and an emission wavelength of 530 nm with a bandwidth of 20 nm.

### Membrane potential measurement

Membrane potential was measured using a BacLight Membrane Potential Kit (Thermo Fisher). Cells were grown in CDM to the desired growth phase. The optical density was then adjusted to an OD_600_ of 0.3 in PBS. DiOC_2_(3) was added to a final concentration of 30 µM, and the bacteria were incubated at 37 °C and gently agitated for 30 min. For plate reader measurements, 200 µl of stained cells was transferred to a Greiner Sensoplate (96-well, black, clear bottom). Using a Tecan Spark plate reader, red and green fluorescence was detected (excitation 488 nm, emission 535 nm and 600 nm), and the red:green fluorescence ratio was calculated. For flow cytometry, stained bacteria were pelleted by centrifugation, resuspended at 1x10^7^ cells/ml in PBS and analysed using a BD FACSCalibur FL-1H and FL-3H.

### Intracellular pH determination

For intracellular pH determination, the fluorogenic probe pHRodo™ Green AM was used. *S. aureus* bacterial cells were grown to the stationary phase, and 0.6 ODU of the bacterial culture was harvested and washed with PBS. The cells were resuspended in 100 µl of pHRodo™ Green AM staining solution (final concentration of 10 µM) and incubated at 37 °C for 30 min. Afterwards, the cells were washed in 300 µl of PBS. Fluorescence was detected in a 96-well Sensoplate (Greiner) using a Tecan Spark plate reader (excitation: 500 nm, emission: 545 nm) or by flow cytometry using a BD FACS Calibur. For calibration of pHRodo Green AM, the “Intracellular pH Calibration Kit” was used according to the manufacturer’s instructions.

### Cell membrane and cell wall costaining

Formaldehyde-fixed *S. aureus* cells were stained with the fixable cell membrane dye FM4-64X (final concentration: 4 µg/ml) and the cell wall dye Bodipy™ Vancomycin-FL conjugate (final concentration: 1 µg/ml) for 10 min on ice. Fluorescence microscopy was performed using a Leica DM5500B fluorescence microscope and the following filter sets (excitation and emission filters): Bodipy™ Vancomycin-FL conjugate: 40-nm BP470 and 50-nm BP525; and FM4-64X: 50-nm BP535 and 75-nm BP610.

### Membrane fluidity measurement

Membrane fluidity was determined via DPH fluorescence polarization. Briefly, one ODU of bacterial cells was harvested by centrifugation, resuspended in prewarmed PBS containing 20 µM DPH (1,6-diphenyl-hexa-1,3,5-triene) and incubated at 37 °C for 15 min. Fluorescence polarization was measured with a BMG CLARIOStar plate reader in a 96-well Sensoplate (Greiner) (excitation: 360 nm-20 nm, emission: 450 nm-10 nm, dichroic filter: LP 410) at 37 °C.

### Antibiotic tolerance assay

To assess the antibiotic tolerance of *S. aureus* HG001 wildtype and (p)ppGpp^0^ cells, either mid-exponential bacteria or late stationary phase bacteria were treated with 100 µg/ml oxacillin (100x MIC of HG001 wildtype and (p)ppGpp^0^) or 50 µg/ml ciprofloxacin (100x MIC of HG001 wildtype and (p)ppGpp^0^). Minimal inhibitory concentrations (MICs) were determined beforehand by the broth microdilution method or E-Test strips in accordance with EUCAST guidelines ^57^. At the indicated time points, CFU counts were performed (as described under “CFU determination” and survival was calculated by comparison to t_0_ (t_treated_/t_0_).

### Statistical analysis

Data were analysed using GraphPad Prism 10 or R. Relevant information regarding statistical tests was added to each figure caption where appropriate. For ANOVA, assumptions of homoscedasticity and normality of the residuals were checked visually by comparing the fitted values with the residuals and comparing the predicted values with the actual residuals. Post hoc tests (posttests) were performed to conduct and correct for multiple comparisons. All tests for which this test was applied were two-tailed. A significance level cut-off of alpha = 0.05 was used. For visual purposes, asterisks denote significance levels as defined in figure captions.

## Data Availability

All data are available from the GEO database under the accession number GSE254567 (https://www.ncbi.nlm.nih.gov/geo/query/acc.cgi?acc=GSE254567).

## Acknowledgements

This work was funded by the Deutsche Forschungsgemeinschaft (Schwerpunktprogramm SPP1879 to CW (Project 423246275)) and by infrastructural funding from the Deutsche Forschungsgemeinschaft (DFG), Cluster of Excellence EXC 2124 ‘Controlling Microbes to Fight Infections’ (Project 390838134).

Sequencing was performed by the Institute for Medical Microbiology (part of the NGS Competence Center NCCT (Tübingen, Germany), while data management, including data storage of the raw data for this project, was performed by the Quantitative Biology Center (QBiC), University of Tübingen, Germany. Mutants from the Nebraska library were obtained through the Network on Antimicrobial Resistance in Staphylococcus aureus (NARSA) program. Figs. 2A, 8A and 9 were created with BioRender.com.

We thank Libera Lo Presti for scientific discussions and editing the manuscript. We acknowledge support from the Open Access Publication Fund of the University of Tübingen.

---

**Fig. S1.**
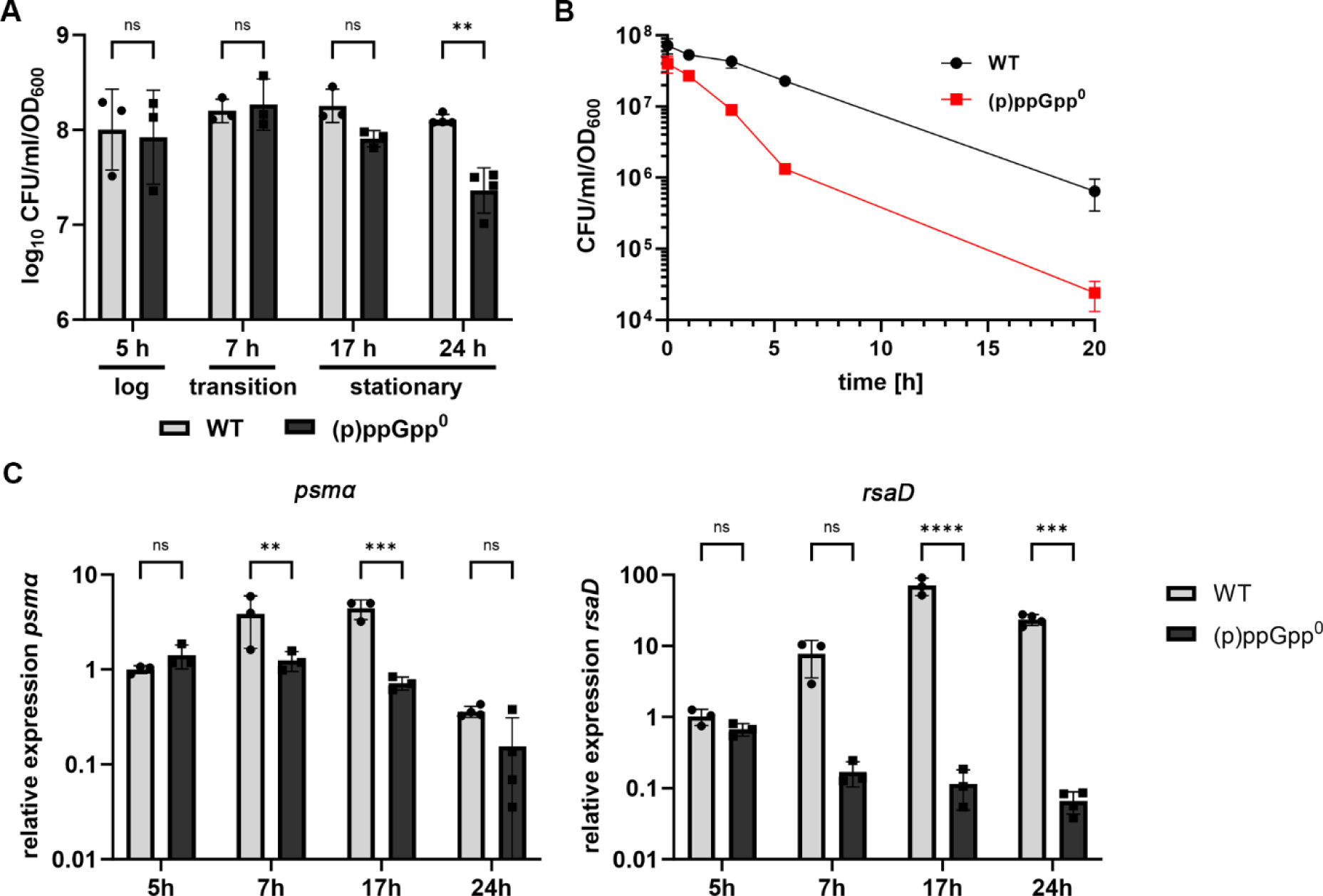
Decreasing cultuarbility of (p)ppGpp^0^ cells during stationary phase and stringent response induction. (A) Culturability of HG001 wildtype (WT) and (p)ppGpp^0^ mutant was evaluated during different growth stages by CFU/ml enumeration normalized to OD_600_. Data shown are mean of ± SD (n=3 biological replicates). Statistical significance was determined by a two-way analysis of variance (ANOVA) with Šidák’s post-test performed on log_10_ transformed data (**p-value <0.01, ns >0.05). (B) Bacteria grown to exponential growth phase were treated with sub-inhibitory concentrations of mupirocin (0,125 µg/ml) and culturability was determined by CFU/ml enumeration normalized to OD_600_. Data shown are mean of ± SD (n=3 biological replicates). (C) Bacterial cells were harvested during different growth phases and total RNA was isolated. *psmα and rsaD* transcript levels were determined by RT-qPCR and normalized to *gyrB* expression by ΔΔCt method. Data are shown as mean ± SD (n=3 biological replicates). Statistical significance was determined by two-way analysis of variance (ANOVA) with Šidák’s post-test (***p-value <0.001, **p-value <0.01, *p-value <0.05, ns >0.05).

**Fig. S2.**
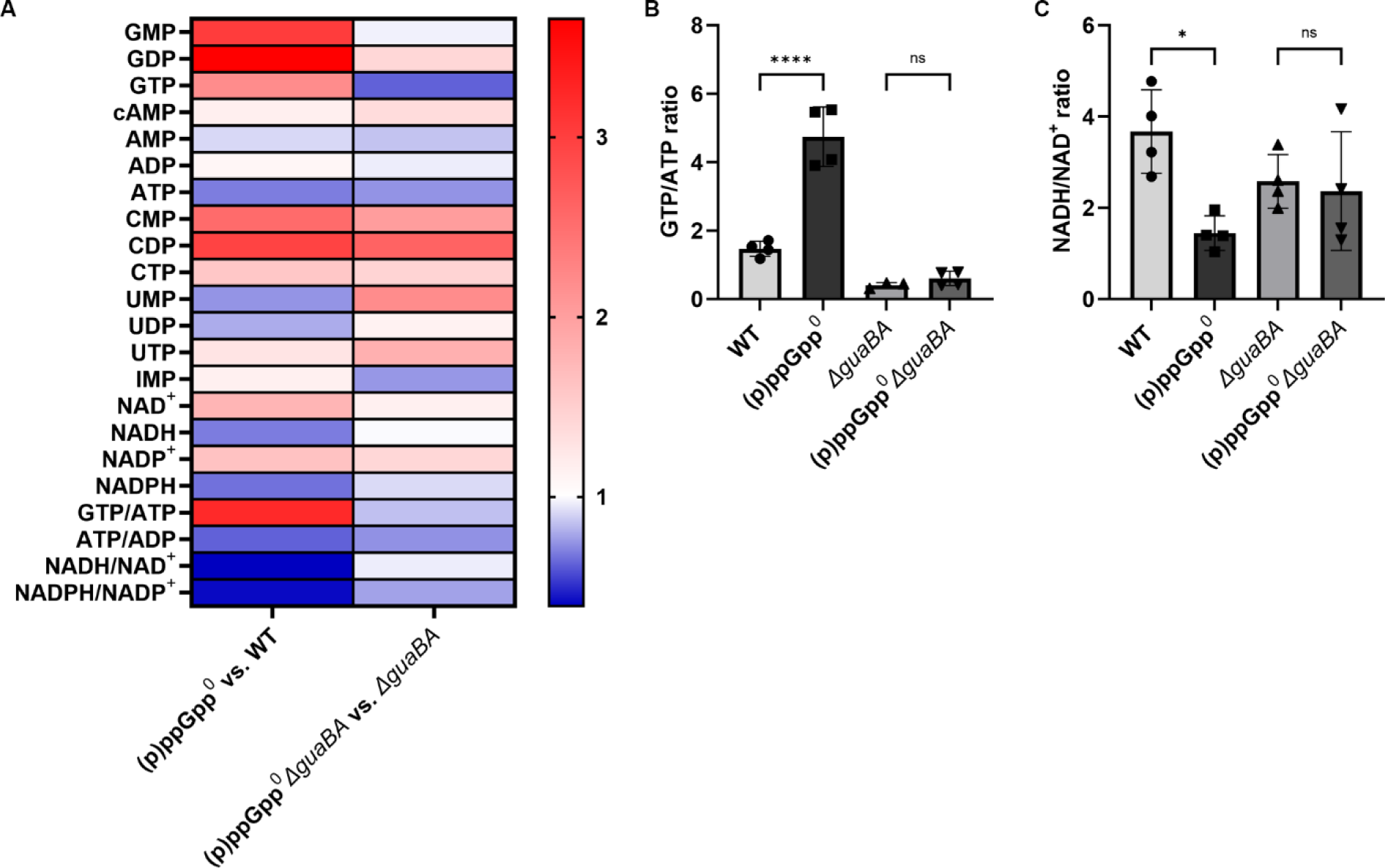
Determination of nucleotide levels in late stationary phase cells. (A) Metabolites were extracted and nucleotides were measured and quantified via LC-MS/MS from cells grown to late stationary growth phase (24h). Shwon are relative nucleotide levels as determined by comparing (p)ppGpp^0^ vs. WT and (p)ppGpp^0^ *ΔguaBA* vs. *ΔguaBA*. (B) The GTP/ATP ratio and (C) the NADH/NAD^+^ ratio were calculated from the relative nucleotide levels. Data shown are mean of ± SD (n=4 biological replicates). Statistical significance was determined by one-way analysis of variance (ANOVA) with Tukey’s post-test (****p-value <0.0001, *p-value <0.05, ns >0.05).

**Fig. S3.**
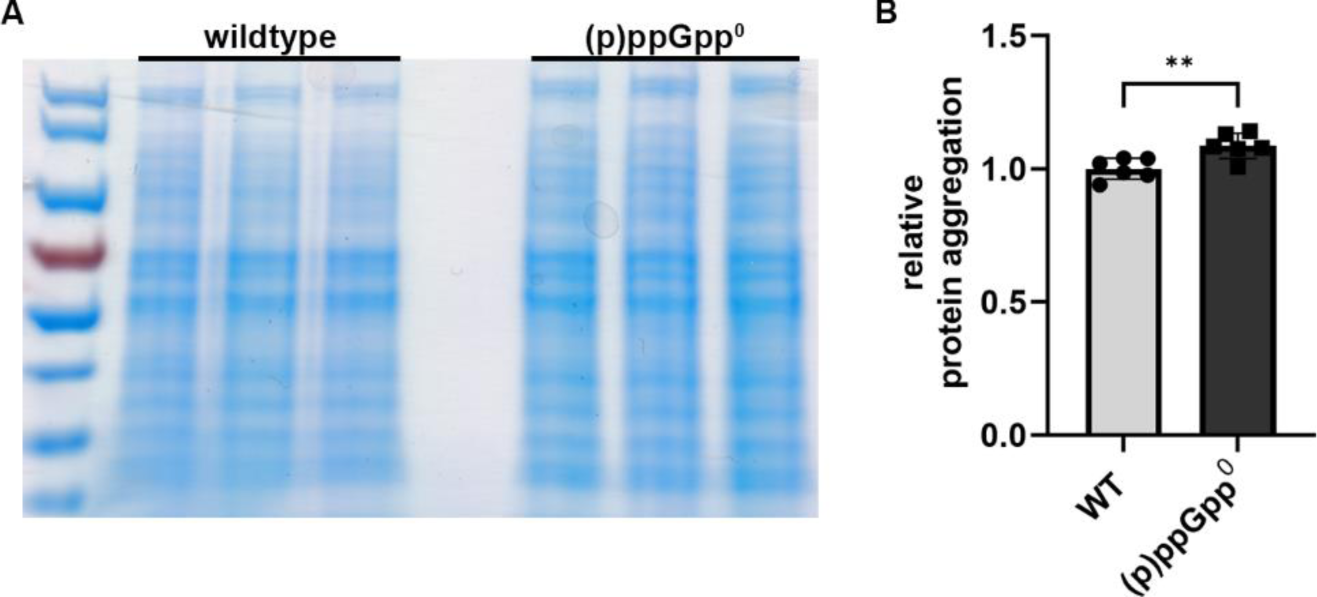
Quantification of protein aggreagates. (A) Protein aggregates were isolated from wildtype and (p)ppGpp^0^ cells grown to late stationary phase and anaylsed by SDS-PAGE and Coomassie staining. For each strain, protein aggregates were isolated from three individual biological replicates (n=3). (B) Protein aggregates were quantified by densiometric analysis. Data shown are mean of ± SD (n=6 biological replicates from two individual experiments). Statistical significance was determined by a two-tailed, unpaired t-test (**p-value <0.01).

**Fig. S4.**
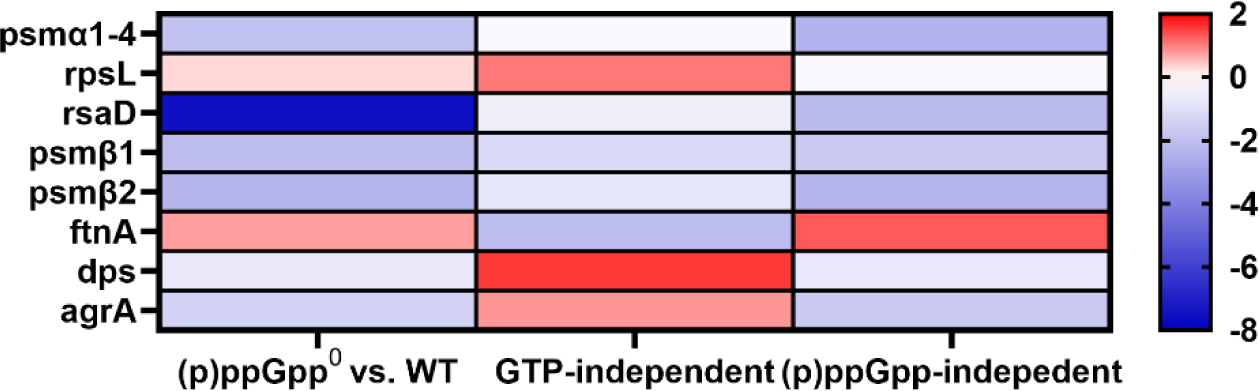
RNA-seq analysis of stringent response genes Heatmaps displaying the RNA-seq fold change of selected stringent response regulated genes. Log2 fold changes in relative transcript abundances are colour-coded with red and blue, indicating up- and downregulation, respectively. “GTP-independent” compares the expression levels of (p)ppGpp^0^ *ΔguaBA* vs. *ΔguaBA*, while “(p)ppGpp-independent” compares the expression levels of (p)ppGpp^0^ vs. (p)ppGpp^0^ *ΔguaBA*. The RNA-seq data shown are from three individual biological replicates (n=3). Full RNA-seq data are available in Table S1 and S2 in the supplementary material.

**Fig. S5.**
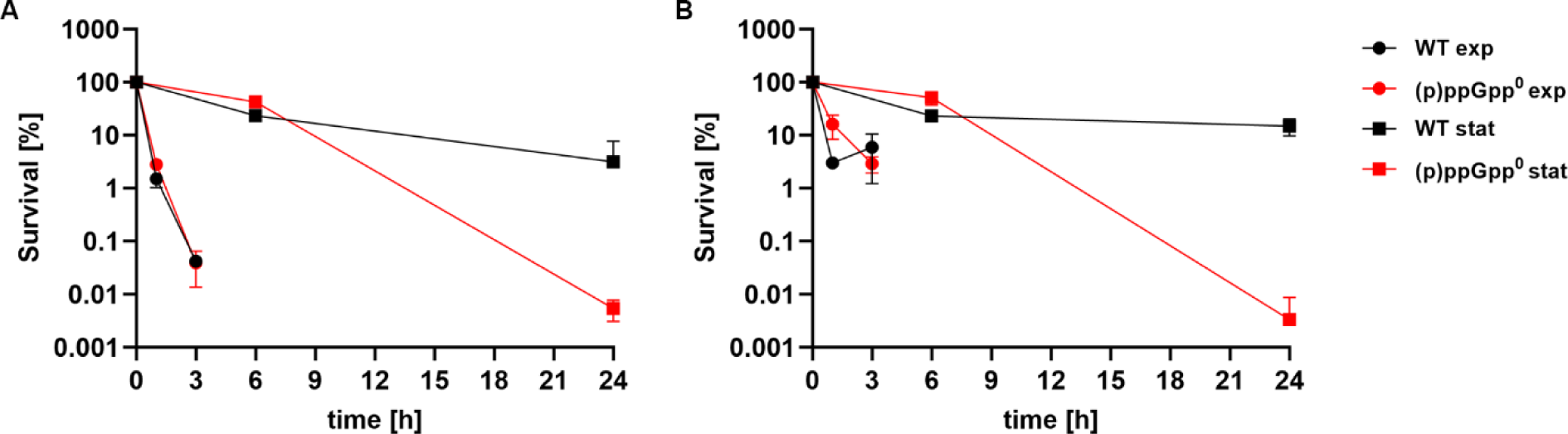
Antibiotic tolerance in exponential and stationary phase. (B) Stationary phase (▪) or mid-exponential phase (●) bacteria were treated with 100x MIC ciprofloxacin (A) or oxacillin (B) and survival was calculated in comparison to untreated cells at different time points after treatment. Data shown are mean of ± SD (n=3 biological replicates).

## References

1 Kaldalu, N. & Tenson, T. Slow growth causes bacterial persistence. Sci Signal 12 (2019). 10.1126/scisignal.aay1167

2. Pontes, M. H. & Groisman, E. A. A Physiological Basis for Nonheritable Antibiotic Resistance. mBio 11 (2020). 10.1128/mBio.00817-20

3. Conlon, B. P., et al. Persister formation in Staphylococcus aureus is associated with ATP depletion. Nat Microbiol 1 (2016). 10.1038/nmicrobiol.2016.51

4 Lempp, M., Lubrano, P., Bange, G. & Link, H. Metabolism of non-growing bacteria. Biol Chem 401, 1479–1485 (2020). 10.1515/hsz-2020-0201

5. Mu, H. et al. Recent functional insights into the magic role of (p)ppGpp in growth control. Comput Struct Biotechnol J 21, 168–175 (2023). 10.1016/j.csbj.2022.11.063

6 Ahn-Horst, T. A., Mille, L. S., Sun, G., Morrison, J. H. & Covert, M. W. An expanded whole-cell model of E. coli links cellular physiology with mechanisms of growth rate control. NPJ Syst Biol Appl 8, 30 (2022). 10.1038/s41540-022-00242-9

7 Potrykus, K., Murphy, H., Philippe, N. & Cashel, M. ppGpp is the major source of growth rate control in E. coli. Environ Microbiol 13, 563–575 (2011). 10.1111/j.1462-2920.2010.02357.x

8 Fernández-Coll, L. & Cashel, M. Possible Roles for Basal Levels of (p)ppGpp: Growth Efficiency Vs. Surviving Stress. Frontiers in Microbiology 11 (2020). 10.3389/fmicb.2020.592718

9 Irving, S. E., Choudhury, N. R. & Corrigan, R. M. The stringent response and physiological roles of (pp)pGpp in bacteria. Nat Rev Microbiol 19, 256–271 (2021). 10.1038/s41579-020-00470-y

10 Hauryliuk, V., Atkinson, G. C., Murakami, K. S., Tenson, T. & Gerdes, K. Recent functional insights into the role of (p)ppGpp in bacterial physiology. Nat Rev Microbiol 13, 298–309 (2015). 10.1038/nrmicro3448

11. Krásný, L., Tišerová, H., Jonák, J., Rejman, D. & Šanderová, H. The identity of the transcription +1 position is crucial for changes in gene expression in response to amino acid starvation in Bacillus subtilis. Molecular Microbiology 69, 42–54 (2008). 10.1111/j.1365-2958.2008.06256.x

12 Geiger, T. et al. The stringent response of Staphylococcus aureus and its impact on survival after phagocytosis through the induction of intracellular PSMs expression. PLoS Pathog 8, e1003016 (2012). 10.1371/journal.ppat.1003016

13 Kriel, A. et al. Direct regulation of GTP homeostasis by (p)ppGpp: a critical component of viability and stress resistance. Mol Cell 48, 231–241 (2012). 10.1016/j.molcel.2012.08.009

14 Kastle, B. et al. rRNA regulation during growth and under stringent conditions in Staphylococcus aureus. Environ Microbiol 17, 4394–4405 (2015). 10.1111/1462-2920.12867

15 Anderson, B. W. et al. The nucleotide messenger (p)ppGpp is an anti-inducer of the purine synthesis transcription regulator PurR in Bacillus. Nucleic Acids Research 50, 847–866 (2021). 10.1093/nar/gkab1281

16 Geiger, T. & Wolz, C. Intersection of the stringent response and the CodY regulon in low GC Gram-positive bacteria. Int J Med Microbiol 304, 150–155 (2014). 10.1016/j.ijmm.2013.11.013

17 Krásný, L. & Gourse, R. L. An alternative strategy for bacterial ribosome synthesis: Bacillus subtilis rRNA transcription regulation. Embo j 23, 4473–4483 (2004). 10.1038/sj.emboj.7600423

18 Carrilero, L. et al. Stringent Response-Mediated Control of GTP Homeostasis Is Required for Long-Term Viability of Staphylococcus aureus. Microbiol Spectr, e0044723 (2023). 10.1128/spectrum.00447-23

19 Salzer, A. & Wolz, C. Role of (p)ppGpp in antibiotic resistance, tolerance, persistence and survival in Firmicutes. Microlife 4, uqad009 (2023). 10.1093/femsml/uqad009

20 Geiger, T., Kastle, B., Gratani, F. L., Goerke, C. & Wolz, C. Two small (p)ppGpp synthases in Staphylococcus aureus mediate tolerance against cell envelope stress conditions. J Bacteriol 196, 894–902 (2014). 10.1128/JB.01201-13

21 Gratani, F. L. et al. Regulation of the opposing (p)ppGpp synthetase and hydrolase activities in a bifunctional RelA/SpoT homologue from Staphylococcus aureus. PLOS Genetics 14, e1007514 (2018). 10.1371/journal.pgen.1007514

22 Horvatek, P. et al. Inducible expression of (pp)pGpp synthetases in Staphylococcus aureus is associated with activation of stress response genes. PLoS Genet 16, e1009282 (2020). 10.1371/journal.pgen.1009282

23 Augagneur, Y. et al. Analysis of the CodY RNome reveals RsaD as a stress-responsive riboregulator of overflow metabolism in Staphylococcus aureus. Molecular Microbiology 113, 309–325 (2020). 10.1111/mmi.14418

24 Sause, W. E. et al. The purine biosynthesis regulator PurR moonlights as a virulence regulator in Staphylococcus aureus. Proc Natl Acad Sci U S A 116, 13563–13572 (2019). 10.1073/pnas.1904280116

25 Goncheva, M. I. et al. Stress-induced inactivation of the Staphylococcus aureus purine biosynthesis repressor leads to hypervirulence. Nat Commun 10, 775 (2019). 10.1038/s41467-019-08724-x

26 Diez, S., Ryu, J., Caban, K., Gonzalez, R. L., Jr. & Dworkin, J. The alarmones (p)ppGpp directly regulate translation initiation during entry into quiescence. Proc Natl Acad Sci U S A 117, 15565–15572 (2020). 10.1073/pnas.1920013117

27 Pu, Y. et al. ATP-Dependent Dynamic Protein Aggregation Regulates Bacterial Dormancy Depth Critical for Antibiotic Tolerance. Mol Cell 73, 143–156.e144 (2019). 10.1016/j.molcel.2018.10.022

28 Fritsch, V. N. et al. The alarmone (p)ppGpp confers tolerance to oxidative stress during the stationary phase by maintenance of redox and iron homeostasis in Staphylococcus aureus. Free Radic Biol Med 161, 351–364 (2020). 10.1016/j.freeradbiomed.2020.10.322

29 Bayer, A. S. et al. Transposon disruption of the complex I NADH oxidoreductase gene (snoD) in Staphylococcus aureus is associated with reduced susceptibility to the microbicidal activity of thrombin-induced platelet microbicidal protein 1. J Bacteriol 188, 211–222 (2006). 10.1128/jb.188.1.211-222.2006

30 Driller, K., Cornejo, F. A. & Turgay, K. (p)ppGpp – an important player during heat shock response. microLife 4 (2023). 10.1093/femsml/uqad017

31 King, A. N. et al. Guanine Limitation Results in CodY-Dependent and -Independent Alteration of Staphylococcus aureus Physiology and Gene Expression. Journal of Bacteriology 200, 10.1128/jb.00136-00118 (2018). doi:10.1128/jb.00136-18

32 Rohmer, C. et al. Influence of Staphylococcus aureus Strain Background on Sa3int Phage Life Cycle Switches. Viruses 14 (2022). 10.3390/v14112471

33 Wang, M., Chan, E. W. C., Wan, Y., Wong, M. H.-y. & Chen, S. Active maintenance of proton motive force mediates starvation-induced bacterial antibiotic tolerance in Escherichia coli. Communications Biology 4, 1068 (2021). 10.1038/s42003-021-02612-1

34 Wan, Y., Chan, E. W. C. & Chen, S. Maintenance and generation of proton motive force are both essential for expression of phenotypic antibiotic tolerance in bacteria. Microbiology Spectrum 0, e00832–00823 (2023). doi:10.1128/spectrum.00832-23

35 Gaca, A. O. et al. Basal levels of (p)ppGpp in Enterococcus faecalis: the magic beyond the stringent response. mBio 4, e00646–00613 (2013). 10.1128/mBio.00646-13

36 Bittner, A. N., Kriel, A. & Wang, J. D. Lowering GTP level increases survival of amino acid starvation but slows growth rate for Bacillus subtilis cells lacking (p)ppGpp. J Bacteriol 196, 2067–2076 (2014). 10.1128/JB.01471-14

37 Pulschen, A. A. et al. The stringent response plays a key role in Bacillus subtilis survival of fatty acid starvation. Mol Microbiol 103, 698–712 (2017). 10.1111/mmi.13582

38 Yang, J., Barra, J. T., Fung, D. K. & Wang, J. D. Bacillus subtilis produces (p)ppGpp in response to the bacteriostatic antibiotic chloramphenicol to prevent its potential bactericidal effect. mLife 1, 101–113 (2022). 10.1002/mlf2.12031

39 Liu, J., Yang, L., Kjellerup, B. V. & Xu, Z. Viable but nonculturable (VBNC) state, an underestimated and controversial microbial survival strategy. Trends Microbiol 31, 1013–1023 (2023). 10.1016/j.tim.2023.04.009

40 Li, J., Liu, C., Wang, S. & Mao, X. Staphylococcus aureus enters viable-but-nonculturable state in response to chitooligosaccharide stress by altering metabolic pattern and transmembrane transport function. Carbohydrate Polymers 330, 121772 (2024). 10.1016/j.carbpol.2023.121772

41 Bai, K. et al. The Role of RelA and SpoT on ppGpp Production, Stress Response, Growth Regulation, and Pathogenicity in Xanthomonas campestris pv. *campestris*. Microbiology Spectrum 9, e02057–02021 (2021). doi:10.1128/spectrum.02057-21

42 Stott, K. V. et al. (p)ppGpp modulates cell size and the initiation of DNA replication in Caulobacter crescentus in response to a block in lipid biosynthesis. Microbiology (Reading) 161, 553–564 (2015). 10.1099/mic.0.000032

43 Turnbough, J., Charles L. Regulation of bacterial gene expression by the NTP substrates of transcription initiation. Molecular Microbiology 69, 10–14 (2008). 10.1111/j.1365-2958.2008.06272.x

44 Mohiuddin, S. G., Hoang, T., Saba, A., Karki, P. & Orman, M. A. Identifying Metabolic Inhibitors to Reduce Bacterial Persistence. Front Microbiol 11, 472 (2020). 10.3389/fmicb.2020.00472

45 Mohiuddin, S. G., Ghosh, S., Kavousi, P. & Orman, M. A. Proton Motive Force Inhibitors Are Detrimental to Methicillin-Resistant Staphylococcus aureus Strains. Microbiology Spectrum 10, e02024–02022 (2022). doi:10.1128/spectrum.02024-22

46 Kaneti, G., Meir, O. & Mor, A. Controlling bacterial infections by inhibiting proton-dependent processes. Biochimica et Biophysica Acta (BBA) - Biomembranes 1858, 995–1003 (2016). 10.1016/j.bbamem.2015.10.022

47 Pohl, K. et al. CodY in Staphylococcus aureus: a regulatory link between metabolism and virulence gene expression. J Bacteriol 191, 2953–2963 (2009). 10.1128/jb.01492-08

48 Bae, T. & Schneewind, O. Allelic replacement in Staphylococcus aureus with inducible counter-selection. Plasmid 55, 58–63 (2006). 10.1016/j.plasmid.2005.05.005

49 Fey, P. D. et al. A genetic resource for rapid and comprehensive phenotype screening of nonessential Staphylococcus aureus genes. mBio 4, e00537–00512 (2013). 10.1128/mBio.00537-12

50 Monk, I. R. & Stinear, T. P. From cloning to mutant in 5 days: rapid allelic exchange in Staphylococcus aureus. Access Microbiol 3, 000193 (2021). 10.1099/acmi.0.000193

51 Monk, I. R., Shah, I. M., Xu, M., Tan, M.-W. & Foster, T. J. Transforming the Untransformable: Application of Direct Transformation To Manipulate Genetically Staphylococcus aureus and Staphylococcus epidermidis. mBio 3, 10.1128/mbio.00277-00211 (2012). doi:10.1128/mbio.00277-11

52 Fuchs, S. et al. AureoWiki ̵ The repository of the Staphylococcus aureus research and annotation community. International Journal of Medical Microbiology 308, 558–568 (2018). 10.1016/j.ijmm.2017.11.011

53 Guder, J. C., Schramm, T., Sander, T. & Link, H. Time-Optimized Isotope Ratio LC–MS/MS for High-Throughput Quantification of Primary Metabolites. Analytical Chemistry 89, 1624–1631 (2017). 10.1021/acs.analchem.6b03731

54 Stahlhut, S. G. et al. The ClpXP protease is dispensable for degradation of unfolded proteins in Staphylococcus aureus. Scientific Reports 7, 11739 (2017). 10.1038/s41598-017-12122-y

55 Maisonneuve, E., Ezraty, B. & Dukan, S. Protein Aggregates: an Aging Factor Involved in Cell Death. Journal of Bacteriology 190, 6070–6075 (2008). doi:10.1128/jb.00736-08

56 George, S. E. et al. Oxidative stress drives the selection of quorum sensing mutants in the Staphylococcus aureus population. Proc Natl Acad Sci U S A 116, 19145–19154 (2019). 10.1073/pnas.1902752116

57. Determination of minimum inhibitory concentrations (MICs) of antibacterial agents by agar dilution. Clinical Microbiology and Infection 6, 509–515 (2000). 10.1046/j.1469-0691.2000.00142.x

