## Supplemental Table S1 - S3 for "(p)ppGpp-mediated GTP homeostasis ensures the survival and antibiotic tolerance of *Staphylococcus aureus* during starvation by preserving the proton motive force"

Table S 1 Strains

| Strain name | Description | Origin |
| --- | --- | --- |
| *Escherichia coli* |  |  |
| IM08B | SA08BΩP_N25_-hsdS (CC8-1) (SAUSA300_0406) of NRS384 integrated between the essQ and cspB genes | (Monk, Tree, Howden, Stinear, & Foster, 2015) |
| *Staphylococcus aureus* |  |  |
| HG001 WT | wildtype, RN1 derivate, *rsbU* repaired | (Herbert et al., 2010; Pohl et al., 2009) |
| HG001 (p)ppGpp^0^ | HG001-229-230-263, Mutation in the synthetase domain of *relP* ,*relQ* and complete deletion of *rel* (ΔrelP_syn_ ΔrelQ_syn_ Δrel) | (Geiger, Kastle, Gratani, Goerke, & Wolz, 2014) |
| USA300 JE2 WT | USA300 derivative, cured of all plasmids | NARSA |
| USA300 JE2 (p)ppGpp^0^ | USA300 JE2 -229-230-263, Mutation in the synthetase domain of *relP* ,*relQ* and complete deletion of *rel* (ΔrelP_syn_ ΔrelQ_syn_ Δrel) | (Horvatek et al., 2020) |
| HG001 (p)ppGpp^0^ *rsh_syn_* compl. | HG001-229-230-263 pCG199 |  |
| HG001 (p)ppGpp^0^ *relP* compl. |  | (Salzer, Keinhorster, Kastle, Kastle, & Wolz, 2020) |
| HG001 (p)ppGpp^0^ *relQ* compl. |  | (Salzer et al., 2020) |
| HG001 *ΔcodY* | HG001-21, *ΔcodY::tetM* | (Pohl et al., 2009) |
| HG001 (p)ppGpp^0^ *ΔcodY* |  | (Salzer et al., 2020) |
| HG001 *ΔpurR* | NE1237, T*nbursa::purR, erm^R^* | This work |
| HG001 (p)ppGpp^0^ *ΔpurR* |  | This work |
| HG001 *ΔguaBA* | HG001-337, markerless *ΔguaBA* mutation | (Kastle et al., 2015) |
| HG001 (p)ppGpp^0^ *ΔguaBA* |  | (Kastle et al., 2015) |
| HG001 WT-PqoxABCDmut | TSS +1 of P*qoxABCD* is mutated from A 🡪 G | This work |
| HG001 (p)ppGpp^0^- PqoxABCDmut | TSS +1 of P*qoxABCD* is mutated from A 🡪 G | This work |
| SH1000 WT | Phage-cured | Susanne Engelmann, TU Braunschweig, Germany |
| SH1000 (p)ppGpp^0^ | SH1000 -229-230-263, Mutation in the synthetase domain of *relP* ,*relQ* and complete deletion of *rel* (ΔrelP_syn_ ΔrelQ_syn_ Δrel) | This work |

Table S2 Plasmids

| Plasmid name | Description | Origin |
| --- | --- | --- |
| pIMAY-Z | Carries Gram-positive ribosome binding site and lacZ cloned downstream from the constitutive cat gene in pIMAY; 8.8 kb, Cm^r^ | (Monk et al., 2015) |
| pCG199 | Integrative *rsh_syn_* complementation plasmid, pCL84-based | (Geiger et al., 2010) |
| pCG216 | Integrative *relQ* complementation plasmid | (Salzer et al., 2020) |
| pCG833 | Integrative *relP* complementation plasmid | (Salzer et al., 2020) |
| pCG919 | pIMAY-Z based plasmid for mutation of TSS+1 P*qoxABCD* is mutated from A 🡪 G | This work |
| pCG229 | pKOR1 with integrated, mutated *relP* | (Geiger et al., 2014) |
| pCG230 | pKOR1 with integrated, mutated *relQ* | (Geiger et al., 2014) |
| pCG263 | pKOR1 with integrated, mutated *rel* | (Geiger et al., 2014) |

Table S3 Ologonucleotides

| Primer name | Sequence 5‘ 🡪 3‘ | Purpose |
| --- | --- | --- |
| psm2391 | CATCGTTTTGTCCTCCTG | qRT-PCR *psmα* |
| psm271 | TCATCGCTGGCATCATTA |  |
| rpslfor | ACCACAAAAACGTGGTGTATGTACT | qRT-PCR *rpsL* |
| rpslseqrev | ACCAGGGATGTATGCGTT |  |
| rsaD-LCfor | GGTAATACACTTGGCTTTTATGGG | qRT-PCR *rsaD* |
| rsaD-LCrev | AGAAGTTATCTCCTTTGTGTTG |  |
| qoxA-LCfor | TCTTTGCTTCTATTATTTGGC | qRT-PCR *qoxA* |
| qoxA-LCrev | GCATGAAGACGATTGAATAAAG |  |
| gyr297 | TTAGTGTGGGAAATTGTCGATAAT | qRT-PCR *gyrB* |
| gyr574 | AGTCTTGTGACAATGCGTTTACA |  |
| pCG919gibfor | AATTCCTGCAGCCCGGGGTTGTTA  TATGGTTCGTCATTTCCA | Insert 1 for cloning of pCG919mut |
| pCG919-2mutfor | CTTAAAATTAATGTTGAGCCCTACA  TTTGTAG |  |
| pCG919-2mutrev | GTAGGGCTCAACATTAATTTTAAGT  TATTACAC | Insert 2 for cloning of pCG919mut |
| pCG919gibrev | GCCGCTCTAGAACTAGTGTCTAGTCAGG  GGCCCCAAC |  |
| pCG919seqfor1 | TTGTTATTCTAACTTCATCTGCAAC | Sequencing primer |
| Pqox-ctrlfor | GAACCCACGTCACACCTTGA |  |
| Tnbuster/ |  | Verification of *purR::Tnbursa* mutant |
| purR-rev | TACGTAACACCACCACTTGC |  |
| relPDIG-for | GTCGCACATTCTTTCAG | verification of *relP* synthase mutant |
| relPDIG-rev | CGTTATTAGGTTTCGTAGAGTT |  |
| relQDIGfor2 | TTCGTAACACTAAAGAAAGTGG | verification of *relQ* synthase mutant |
| relQDIGrev2 | GCGTGTAATATTTTTGAGCT |  |
| rel431for | GCGTGGCTTTATCATTGG | verification of *rel_syn_* mutant |
| relLC4rev | ACTTCAACCATCATTCGG |  |

Geiger, T., Goerke, C., Fritz, M., Schafer, T., Ohlsen, K., Liebeke, M., . . . Wolz, C. (2010). Role of the (p)ppGpp synthase RSH, a RelA/SpoT homolog, in stringent response and virulence of Staphylococcus aureus. *Infect Immun, 78*(5), 1873-1883. doi:10.1128/IAI.01439-09

Geiger, T., Kastle, B., Gratani, F. L., Goerke, C., & Wolz, C. (2014). Two small (p)ppGpp synthases in Staphylococcus aureus mediate tolerance against cell envelope stress conditions. *J Bacteriol, 196*(4), 894-902. doi:10.1128/JB.01201-13

Herbert, S., Ziebandt, A. K., Ohlsen, K., Schäfer, T., Hecker, M., Albrecht, D., . . . Götz, F. (2010). Repair of global regulators in Staphylococcus aureus 8325 and comparative analysis with other clinical isolates. *Infect Immun, 78*(6), 2877-2889. doi:10.1128/iai.00088-10

Horvatek, P., Salzer, A., Hanna, A. M. F., Gratani, F. L., Keinhorster, D., Korn, N., . . . Wolz, C. (2020). Inducible expression of (pp)pGpp synthetases in Staphylococcus aureus is associated with activation of stress response genes. *PLoS Genet, 16*(12), e1009282. doi:10.1371/journal.pgen.1009282

Kastle, B., Geiger, T., Gratani, F. L., Reisinger, R., Goerke, C., Borisova, M., . . . Wolz, C. (2015). rRNA regulation during growth and under stringent conditions in Staphylococcus aureus. *Environ Microbiol, 17*(11), 4394-4405. doi:10.1111/1462-2920.12867

Monk, I. R., Tree, J. J., Howden, B. P., Stinear, T. P., & Foster, T. J. (2015). Complete Bypass of Restriction Systems for Major Staphylococcus aureus Lineages. *mBio, 6*(3), e00308-00315. doi:10.1128/mBio.00308-15

Pohl, K., Francois, P., Stenz, L., Schlink, F., Geiger, T., Herbert, S., . . . Wolz, C. (2009). CodY in Staphylococcus aureus: a regulatory link between metabolism and virulence gene expression. *J Bacteriol, 191*(9), 2953-2963. doi:10.1128/jb.01492-08

Salzer, A., Keinhorster, D., Kastle, C., Kastle, B., & Wolz, C. (2020). Small Alarmone Synthetases RelP and RelQ of Staphylococcus aureus Are Involved in Biofilm Formation and Maintenance Under Cell Wall Stress Conditions. *Front Microbiol, 11*, 575882. doi:10.3389/fmicb.2020.575882
